# Imaging the dynamics of uterine contractions in early pregnancy

**DOI:** 10.1101/2023.12.06.570447

**Authors:** Madeline Dawson, Diana Flores, Lisa Zou, Shivani Anandasenthil, Rohit Mahesh, Olmo Zavala, Ripla Arora

## Abstract

The myometrium or smooth muscle of the uterus contracts throughout the life of the organ. Uterine muscle contractility is essential for reproductive processes including sperm and embryo transport, and during the uterine cycle to remove menstrual effluent or estrus debris. Even still, uterine contractions have primarily only been studied in the context of preterm labor. This is partly due to a lack of methods for studying the contractile characteristics of the uterine muscle in the intact organ. Here, we describe an imaging-based method to evaluate the contractility of both the longitudinal and circular muscles of the uterus in the cycling stages and in early pregnancy. By transforming the image-based data into 3D spatiotemporal contractility maps, we calculate waveform characteristics of muscle contractions, including amplitude, frequency, wavelength, and velocity. We report that the native organ is highly contractile during the progesterone-dominant diestrus stage of the cycle when compared to the estrogen-dominant proestrus and estrus stages. We also observed correlations between contractility during pre-implantation stages of pregnancy and observed embryo movement patterns. During the first phase of embryo movement when clustered embryos move towards the middle of the uterine horn, uterine contractions are dynamic and non-uniform between different segments of the uterine horn. In the second phase of embryo movement, contractions are more uniform and rhythmic throughout the uterine horn. Finally, when our method is applied to *Lpar3* mutant uteri that display faster embryo movement, we observe global and regional increases in contractility. Our method provides a means to understand the wave characteristics of uterine smooth muscle in response to modulators and in genetic mutants. Better understanding uterine contractility in the early pregnancy stages is critical for the advancement of artificial reproductive technologies and a possibility of modulating embryo movement during clinical embryo transfers.

## INTRODUCTION

Smooth muscle performs organ-specific function throughout the body (Webb, 2003). For instance, smooth muscle in the stomach and intestines aids in food peristalsis and nutrient absorption (Hafen and Burns, 2023) while smooth muscle in the bladder wall is responsible for the whole organ contraction and relaxation during retention and urination (Andersson and Arner, 2004). In regard to the reproductive tract, uterine smooth muscle plays an important role in the movement of eggs to the fundus during ovulation, or sperm towards the fallopian tube (oviduct) during fertilization. However, contractions are also key to a woman’s non-pregnant state in discarding menstrual effluent and removing eggs released every cycle. Efforts have been made towards measuring uterine contractions in late-term pregnancy due to the challenges caused by preterm labor. Contractions in the non-pregnant but cycling uterus and the early pregnant uterus differ from the late-term pregnant organ (Aguilar and Mitchell, 2010). Thus, imaging methodologies must be designed to capture contractile activity in the non-pregnant and early pregnant uterus.

In all mammals, the uterine smooth muscle is derived from the Mullerian duct (embryonic precursor of the uterus) mesenchyme (Massman and Harland, 1980). In rodents (rats and mice), each Mullerian duct develops into a uterine horn, whereas the two horns fuse to form a single uterine chamber in primates (humans). In both mice and humans, the uterine muscle surrounds the inner epithelium (endometrium) and the mesenchyme (stroma) and comprises of an outer layer of longitudinal muscle and an inner layer of circular muscle. While in most mammals there is a clear separation between the longitudinal and circular muscle layers, this separation is less clear in larger mammals such as humans (Massman and Harland, 1980). Instead, there exists a junctional muscle zone (displays properties of circular muscle) that separates the outer uterine muscle (displays properties of longitudinal muscle) from the inner endometrium (Aguilar and Mitchell, 2010). Despite the two distinct muscle layers in mice, a junctional muscle zone was recently reported supporting the utility of mouse as a model to understand the patterns of and the mechanisms regulating uterine contractions (Kagami et al., 2020).

Uterine contractions can be measured using invasive techniques such as intrauterine pressure measurement (Künzel et al., 2014; Robuck et al., 2018) and electromyography (Bulletti and de Ziegler, 2006) or non-invasive molecular imaging techniques such as 3D ultrasounds (Bulletti and de Ziegler, 2006) or magnetic resonance imaging (Lam et al., 2018). In vivo, pressure recordings and ultrasound can themselves contribute to contractility and MRI and ultrasounds do not give spatial and quantitative waveform metrics such as amplitude, frequency, and velocity (Lovasz et al., 2013; Qu et al., 2021; Ter Haar et al., 1978). Uterine contractions have been measured by isolating either tissue strips from the uterus (Allix et al., 2008; Bafor et al., 2017; Dodds et al., 2015; Smith et al., 2007) or transverse uterine slices (Qu et al., 2021). Tissue pieces are equilibrated in an organ bath (Talo, 1980) and spontaneous contractions or the effect of agonist induced contractions is estimated. It is critical to note that often, in these methods, spontaneous contractions are measured after a mechanical force is applied using a tension transducer. Limitations of these methods include variability in data generated as different transducer forces will result in different contractile activity which may not reflect spontaneous contractility. Further separating the uterine muscle from the endometrium or slicing the uterus leads to loss of spatial information (Mackler et al., 1999).

During the estrus cycle, using uterine strips from rat uteri in an organ bath under the effect of a tension transducer, electric field-induced contractility was highest at the estrus stage, and the lowest responses were observed at the diestrus stage (Houdeau et al., 2003). In vivo imaging of the contractions of the rat uteri was attempted by Crane and Martin (Crane and Martin, 1991a) and generated contrasting results. In this study, three methods were used: a balloon was inserted into the lumen to study intraluminal pressure, electrodes were attached to both the cervical and oviductal ends of the uterus to measure contractions, and laparoscopy was used on anesthetized rats to record the uterus moving in vivo. The use of balloon and electrodes caused rats to lose their cyclicity and thus laparoscopy surgery was used to evaluate contractions during estrus cycle. Qualitatively, total contractile activity was determined to be lowest at proestrus, increased at estrus, and highest during diestrus. The entire mouse uterine horn has also been subjected to a video recording in the non-pregnant state in vitro (Dodds et al., 2015). Similar to the rat uterus in vivo recordings, this method showed that even in the mouse, diestrus is the most contractile stage. However, when tension transducer was applied to the intact uterine horn and contractions were quantified under mechanical tension, diestrus showed low frequency and low amplitude contractions and proestrus showed high frequency and high amplitude contractions. These studies highlight conflicting data on which stage of the oestrus cycle is most contractile.

An advancement of the whole horn method (Dodds et al., 2021) used pressure readings to calculate amplitude, image-based changes in uterine horn edges to calculate contraction velocity, frequency and directionality and electrical-induced activity to measure spike potential and burst duration for a proestrus staged uterus. This method and others have shown that fluid mediated distension increases uterine contractility (Dodds et al., 2021; Hellman et al., 2018). Most recently, imaging-based methods have been used to measure both strength and directionality of contractions in the proestrus stage of the mouse uterus using in vivo recordings and spatiotemporal mapping (Zhang et al., 2019). The limitations of this study are that only proestrus stage, which displays limited uterine length and lower strength contractions, could be recorded. Further, the use of anesthetics could potentially influence smooth muscle contractions. None of the reported methods permit the study of early stages of pregnancy to evaluate the relationship between uterine contractility and early pregnancy embryo movement.

In all prior studies that used spatiotemporal mapping to examine contractility, the change in diameter between uterine edges was used as a proxy for muscle contraction. Thus, these methods quantify contractions caused by the circular smooth muscle but do not account for longitudinal muscle contractions (Dodds et al., 2015). On the other hand, when isolating uterine strips, the contractility would depend on the direction in which the strips are loaded into the bath and where the tension transducers are applied to determine whether circular or longitudinal muscle activity was measured (Houdeau et al., 2003). All methods to date fail to report combined contributions of the circular and longitudinal smooth muscle to uterine contractility.

Contractions are critical to embryo movement in the uterus (Flores et al., 2020), there is an urgent need to develop methods to evaluate uterine contractility during early pregnancy. For pregnant mice, oviductal contractions have been quantified by recording changes in the oviductal wall and tracing the movement of the egg/embryo, and directly correlating the movement of the muscle wall with the movement of the egg in the oviductal space (Dixon et al., 2009). Studies such as these can’t be performed in the uterus as the thickness of the murine smooth muscle layer (∼500μm) prevents optical light from passing through and reaching the uterine lumen where the embryos are residing. Optical coherence tomography has been used to track sperm and oocyte movement in the oviduct (Wang and Larina, 2023), however, this methodology has not been successfully applied to the uterus yet. Here, we discuss an ex vivo imaging-based method to trace uterine contractions during early pregnancy in tomato transgene expressing reporter mice. First, we apply this method to different stages of the estrus cycle and compare our results to previously published data. We then apply our method to pre-implantation stages of pregnancy and compare contractility patterns to our recently published embryo movement patterns (Flores et al., 2020). Finally, we apply this method to genetic mouse models with disrupted embryo movement patterns (*Lpar3^-/-^*)(Flores et al., 2020; Hama et al., 2007; Ye et al., 2005) and detect global and spatial differences in uterine contractility that explain the differential embryo movement patterns.

## METHODS

### Animals

All animal research was carried out under the guidelines of the Michigan State University Institutional Animal Care and Use Committee. CD1 (ICR), wildtype C57BL/6J, and *Lpar3^tm1JCh^*(*Lpar3^+/-^* and *Lpar3^-/-^*) mice (Ye et al., 2005) carrying the Rosa mTmG allele (Muzumdar et al., 2007) were generated. Mice were maintained on a 12 h light/dark cycle. For nonpregnant studies, the estrus stage was determined using vaginal smear cytology (McLean et al., 2012). Dissections were performed at each stage of estrus: estrus, proestrus, metestrus, and diestrus. For pregnancy studies, adult females aged 6 to 12 weeks, were mated with fertile wildtype males to induce pregnancy. The appearance of a vaginal plug was identified as GD0.5 or GD0 1200h. For CD1 females, uterine dissections were performed on times when embryos are moving in the uterus - GD3 at 0600 h, 1200 h, and 1800 h and post-implantation - GD4 at 1200 h. In wildtype, *Lpar3*^+/−^ and *Lpar3^-/-^* (C57BL/6J background), dissections were performed on GD3 at 0600 h, 1200 h, and 1800 h and on GD4 at 1200 h. A minimum of three mice were analyzed for each condition to ensure data reproducibility in independent events.

### Ex-vivo, spatiotemporal video recording of the contracting uterus

Immediately after sacrificing the mouse, the uterus was harvested and secured in a plastic petri dish using syringe needles. Three needles were used to hold the uterus in place within the petri dish (one pierced through each ovary and one pierced through the cervix) to mimic the position of the uterus in vivo (**Fig. 1A**). The petri dish containing a pinned uterus immersed in Phosphate Buffer Saline (PBS) was then placed under a Leica MZ10F fluorescence stereo microscope with a fluorescence filter for Texas Red. Time-lapse uterine contractions were recorded using the LASX software with images captured every 200msec for 5 minutes (total 1500 frames). After this initial video was taken, 100 μl of 10mg/ml solution of salbutamol sulfate prepared in 1:9 Ethanol:PBS, was added uniformly to stop the uterus from contracting (Flores *et* al, 2020). Another set of images was acquired for 5 minutes as a non-contracting control. Next, .lif files from the LASX software were converted to .mp4 files and edited using the virtual dub application (ver 1.9.11). Virtual dub was used to crop .mp4 videos, to get one uterine horn per video, and to threshold tomato intensity and increase the contrast between the uterus and the video background (**Fig. 1B**). Note that thresholding and changing contrast doesn’t change the relative changes in tomato intensity across the entire uterine horn. Care should be taken to not saturate the tomato intensity as contractile differences rely on changes in tomato intensity along the uterine horn. .mp4 videos were then converted into .avi files using the ‘Xvid MPEG-4 codec’ compression option within virtual dub. Each video produced two .avi files, one for each uterine horn (**Supplementary Movie 1**). These .avi files were then subjected to image analysis to calculate contractility waveform metrics.

**Figure 1:**
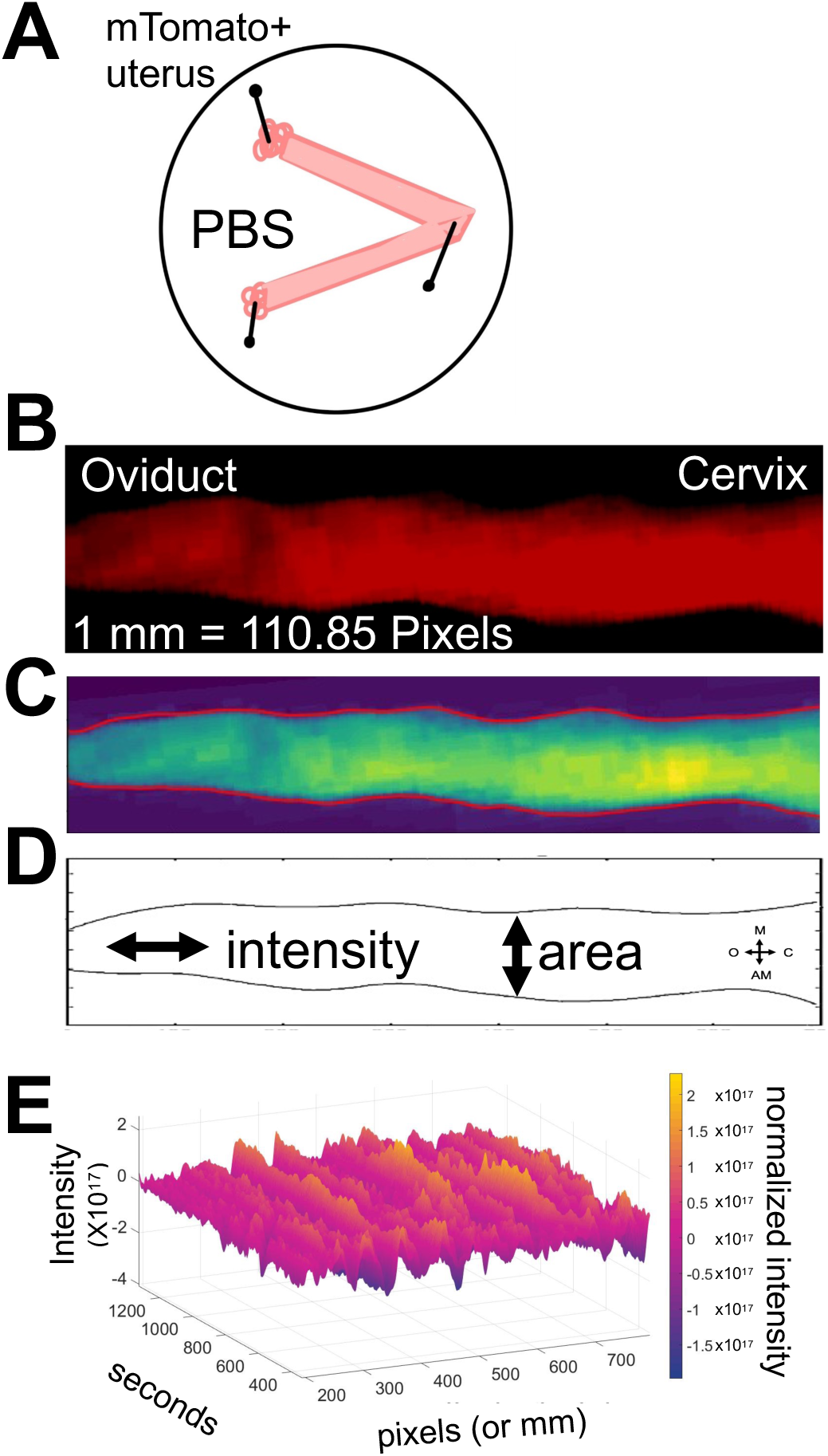
Set up for uterine horn ex vivo imaging and image analysis. (A) Schematic of how the uterine horns are pinned after dissection. (B) Example frame from uterine contraction video displaying tomato positive uterine horn. (C) Smoothened image of (B) showing variation in contraction intensity with blue being the most relaxed and yellow being the most contracting region. (D) schematic of the two types of measurements made for evaluating contractions. (E) Example 3D graphs obtained from image processing with X axis signifying uterine horn distance in pixels or mm, Y axis signifying frames or time in seconds and Z axis in color signifying contraction either as changes in area or changes in tomato intensity.

### Methodology for extracting waveform metrics from uterine contraction videos

Our methodology aims to identify and characterize two types of contractions observed in videos of the uterine horn (**Fig. 1B, C**). The first type of contraction, believed to arise from the circular muscle, is discerned via changes in the uterine horn’s area within the plane perpendicular to the camera’s view (**Fig. 1D**). The second contraction type, hypothesized to originate from both the longitudinal and circular muscle, is detected through alterations in pixel intensity within the video frames (**Fig. 1D**). These intensity changes correlate with dye-density shifts and the number of cells intersected by planes parallel to the camera’s viewpoint.

Our methodology contains three main steps: identifying the uterine horn in each frame of the videos, estimating the changes in area and intensity through time at each longitudinal position of the uterine horn, and characterizing the contractions detected by estimating the amplitude, frequency, velocity and wavelength of the contractions.

#### Step 1: Uterine Horn Identification

The initial step in video preprocessing involves automatic detection of the uterine horn’s upper and lower boundaries in each frame and column. This task is challenging in areas where the uterine horn is exceedingly thin, as the camera sensor yields minimal intensity gradients at the horn’s boundary, making precise boundary determination difficult. Additionally, blood stains in the petri dish introduces further intensity gradients that may confound the detection of the horn’s start and end points.

To delineate the horn’s boundaries (**Supplementary Movie 2**), we first apply a horizontal and temporal smoothing process to the videos (**Fig. 2A**), which mitigates any noise artifacts that might interfere with automatic detection. This smoothing is accomplished using a 3-size Gaussian filter in time and a 5-size Gaussian filter in space, implemented via the GaussianBlur function of OpenCV (Bradski and Kaehler, 2008). These filter parameters were manually selected based on a subjective comparison of the original and smoothed video frames. Importantly, this video intensity smoothing is solely for accurate uterine horn boundary selection and is not utilized in subsequent intensity analyses.

**Figure 2:**
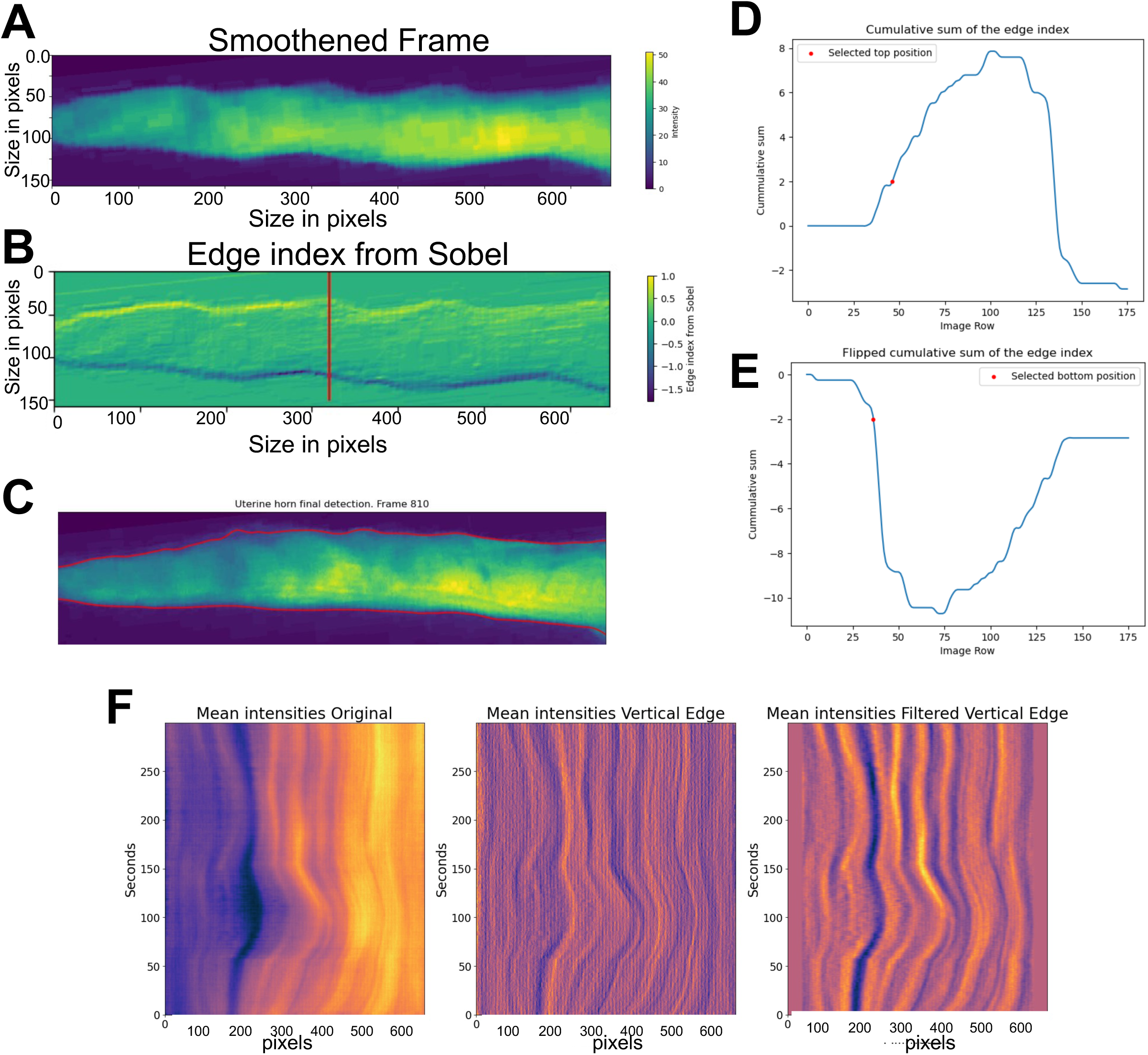
Extracting area and intensity information from contraction videos. (A) Smoothened image of one frame of an example uterine contraction video frame. (B) An example of the locations obtained from the edge index image and the top and bottom borders of the image (C) together with the cumulative distribution (D) and the flipped cumulative distributions (E) of the middle column in B. (F) Intensity data obtained from image segmentation and example of how data is normalized to uterine horn area.

Next, we employ the Sobel filter to detect horizontal and vertical edges within each video frame. The Sobel filter’s size in both directions is 5×5. To give greater weight to vertical edges and generate a single edge index image, vertical edges are weighted according to the formula:

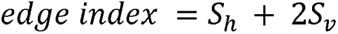

where *S_h_* is the horizontal Sobel filter and *S_v_* is the vertical Sobel filter. **Figure 2B** illustrates an example of the edge index derived from smoothing a single frame and employing Sobel to primarily detect vertical edges.

From the *edge index* image, the top position is identified for each column as the first row-wise location that exceeds two from the cumulative *edge index* of the column (**Fig. 2D**). The bottom position is identified as the first row-wise location that is lower than two from the flipped cumulative sum of the *edge index* of the column (**Fig. 2E**). These two parameters are the ones that worked the best for the analyzed videos (**Fig. 2C**). As long as the videos have similar intensities, the detected start and end row locations of the uterus horn are not very sensitive to selecting these parameters. The final positions get further smoothened by cubic interpolation. For both positions, top and bottom, the total number of knot points used is one every ten columns.

#### Step 2. Area and Intensity Change Estimation

The locations established in Step 1 serve to extract initial and final positions, as well as mean intensity values for each column in each frame of the analyzed videos. We then utilize these values to examine the temporal behavior of the two muscle types via Hovmoller plots (Ernest, 1949). These plots feature the spatially analyzed value (area or mean intensity) on the x-axis and time on the y-axis. To identify space changes, we employ a 5-size Sobel kernel filter, striking a balance between filtering minor spatial and temporal variations and detecting changes in each field. Finally, we filter the low frequency signal in space and then apply a low-pass filter to remove noise again. **Figure 2F** shows the result of this step of a complete video for the intensity variable. For further details see supplementary methods.

#### Step 3. Contraction Characterization

Intensity or area based output plots (**Fig. 3A**) were used to quantify uterine horn contractions. We compared 3D plots from a contracting uterus (**Fig. 3A**) and corresponding salbutamol treated non-contracting uterus (**Fig. 3A’**) to ensure that changes observed were indicative of muscle movement and not changes in uterine shape along the length of the horn. Contraction characterization is achieved using a technique commonly employed in meteorology, where we identify lines in the Hovmoller plot (Ernest, 1949) and correlate these with the properties of a moving wave. Amplitude and period were calculated as from the distance (pixels)-time (frames) graph. Time was converted from frames to seconds, with 200 frames per second resulting in 1500 frames or 300 seconds. Distance is reported in pixels for this study where 110.85pixels = 1mm. Frequency was calculated as the inverse of period. The wave’s velocity is determined by the line’s slope, with flatter lines indicating faster speeds. The wave’s direction is dictated by the slope of the line, with positive and negative slopes signifying rightward (towards cervix) and leftward (towards oviduct) movement, respectively. Using the slope of the line we did observe waves traveling towards both the oviduct and the cervix, however for this study, we focused on evaluating the spatial distribution of contractions and did not focus on the directionality of contractions. Wavelength is calculated as velocity divided by frequency. For segment analysis, each ‘mean intensities’ graph was sectioned into thirds using lines perpendicular to the x-axis at each third of the x-axis, effectively splitting the graph into oviductal, middle and cervical segments (**Fig.3B**). Three waves were selected at random from each segment and wave function analysis was performed on a total of 9 waves per uterine horn. These wave metrics were then used to compare contractions between different experimental groups.

**Figure 3:**
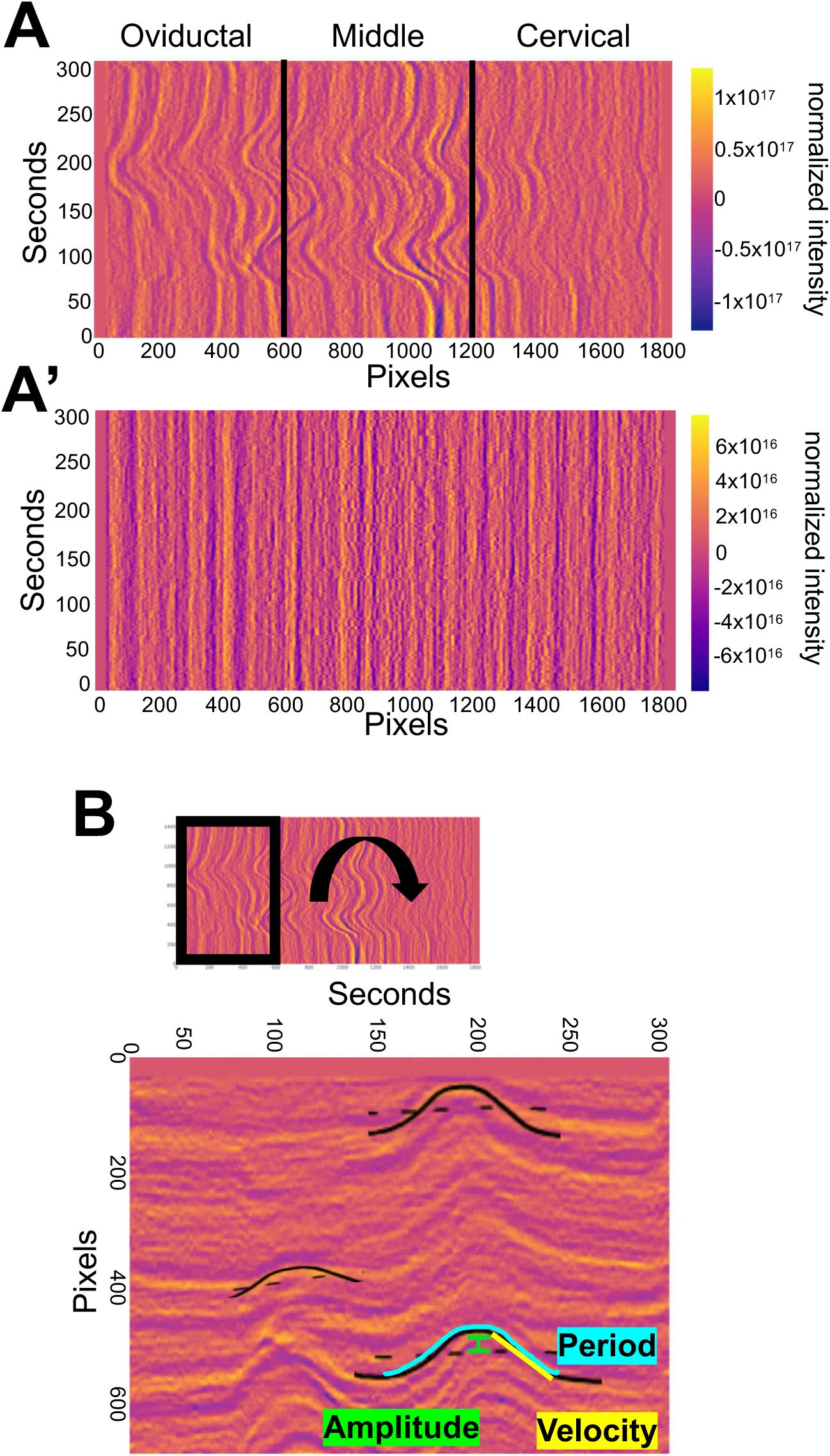
Calculating waveform metrics. (A) Example 3D image of mean intensities showing contractile wave activity in three segments of the uterine horn – oviductal, middle and cervical. (A’) Uterine horn in A treated with salbutamol showing no contractile activity. (B) Example of how waveform metrics such as amplitude, velocity and period are calculated from the waves in the 3D plots.

### Whole-mount immunofluorescence

Whole-mount immunofluorescence staining for wildtype, *Lpar3^+/-^*and *Lpar3^-/-^* uteri was performed as described previously (Arora et al., 2016). Uteri were fixed in DMSO:Methanol (1:4) after dissection. To stain the uteri, they were rehydrated for 15 minutes in 1:1, Methanol: PBST (PBS, 1% Triton X-100) solution, followed by a 15 minutes wash in 100% PBST solution before incubation. Samples were incubated with Hoechst (B2261, Sigma-Aldrich, St. Louis, MO, USA) diluted in PBST (1:500) for two nights at 4°C. The uteri were then washed once for 15 minutes and three times for 45 minutes each, using PBST. Next, the uteri were stretched in 100% methanol, followed by 30 minutes dehydration in 100% methanol, an overnight incubation in 3% H2O2 solution diluted in methanol, and a final dehydration step for 60 minutes in 100% methanol. Finally, samples were cleared using a 1:2 mixture of Benzyl Alcohol: Benzyl Benzoate (108006, B6630, Sigma-Aldrich, St. Louis, MO, USA).

### Confocal microscopy

Confocal imagining procedures were done as previously described (Flores et al., 2020). Stained uteri were imaged using a Leica TCS SP8 X Confocal Laser Scanning Microscope System with white-light laser, using a 10x air objective. For each uterine horn, z-stacks were generated with a 7.0 µm increment, and tiled scans were set up to image the entire length and depth of the uterine horn (Arora et al., 2016). Images were merged using Leica software LASX version 3.5.5.

### Image analysis for embryo location

Image analysis was done using commercial software Imaris v9.2.1 (Bitplane, Zurich, Switzerland). Embryo location was assessed as described previously (Flores et al., 2020). Briefly, confocal LIF files were imported into the Surpass mode of Imaris and Surface module 3D renderings were used to create structures for the oviductal-uterine junctions, embryos, and horns. The three-dimensional Cartesian coordinates of each surface’s center were identified and stored using the measurement module. The distance between the oviductal-uterine junction and an embryo (OE), the distance between adjacent embryos (EE), and the horn length was calculated using the orthogonal projection onto the XY plane. All distances were normalized to the length of their respective uterine horn. Horns with less than three embryos were excluded from the analysis. These distances were used to map the location of the embryos relative to the length of the uterine horn. The uterine horn was divided into three equally spaced segments – closest to the oviduct, middle, and closest to the cervix. These segments were quantified for the percentage of embryos present in each section. Embryos in the oviductal region close to the oviductal-uterine junction were accounted for in the first oviductal segment.

### Statistical Analysis

Data groups were compared using the Mann-Whitney test (unpaired experimental design, nonparametric test, compare ranks) with resulting two-tailed p-values. P-values <0.05 were considered significant. Statistical Analysis was performed using GraphPad Prism 8.2.1.

Outliers were identified and removed using the ROUT (robust regression and outlier) method (Motulsky and Brown, 2006) with Q, the maximum desired false discovery rate, set at 1%. The ROUT method uses n outlier detection method, based on the false discovery rate, to choose outliers that are outside the prediction of the model.

## RESULTS

### Ex vivo method to measure and compare uterine contractility

Typically, uterine horns vary in length based on genetic background (mixed background CD1/ICR mice have longer uterine horns than C57Bl6 mice). Further, during pregnancy, there is proliferation and increase in the length of the uterine tissue (Finn, 1968). Thus, it is key to measure contractility in a dynamic range of uterine horn lengths and different pregnant and non-pregnant stages. Our method preserves the structure of the uterine horn and relative orientation in the in vivo setting. In vivo, the ovaries are attached to the kidneys (fixed structures) and the cervix is attached to the body wall. We mimic this configuration in the petri dish by pinning the oviductal and cervical ends while allowing spontaneous contractility to occur freely in the body of the uterus.

### Measurement of contractile behavior using change in area vs change in tomato intensity

Contractility can be measured based on the movement of the edges of the smooth muscle that surrounds an epithelial lumen. This method has been employed to measure contractility in the oviduct (Bianchi et al., 2021; Dixon et al., 2009) and the non-pregnant uterus (Dodds et al., 2021; Zhang et al., 2019). Further, since the uterine cells express tomato reporter, increase in tomato intensity highlights a contracting region while loss of tomato intensity suggests muscle relaxation. Uterine smooth muscle is comprised of an outer longitudinal layer and an inner circular layer. Considering that we are using whole tissue 2D images for estimating contractility, changes in tomato intensity likely reflect combined effects from both longitudinal and underlying circular smooth muscle layers (**Fig. 1-3**).

We first compared data generated from the 3D area plots to the 3D intensity plots across the proestrus and diestrus stage (**Fig. 4A-D**). The 3D plots suggested that area plots often had regions where contractility was not apparent although in the corresponding regions of the intensity plots, contractility was observed. We wanted to spatially analyze contractility during preimplantation stages, thus we used contractility measurements originating from the intensity plots and not the area plots. When comparing values of different waveform metrics, we observed that the range of amplitude, velocity, frequency and wavelength were relatively similar across both area and intensity plots. We also observed that all of these metrics were much higher at the diestrus stage (**Fig. 4B, D**) compared to the proestrus stage (**Fig. 4A, C**). These data suggest that spontaneous contractions are stronger and more frequent in the diestrus stage when compared to the proestrus stage.

**Figure 4:**
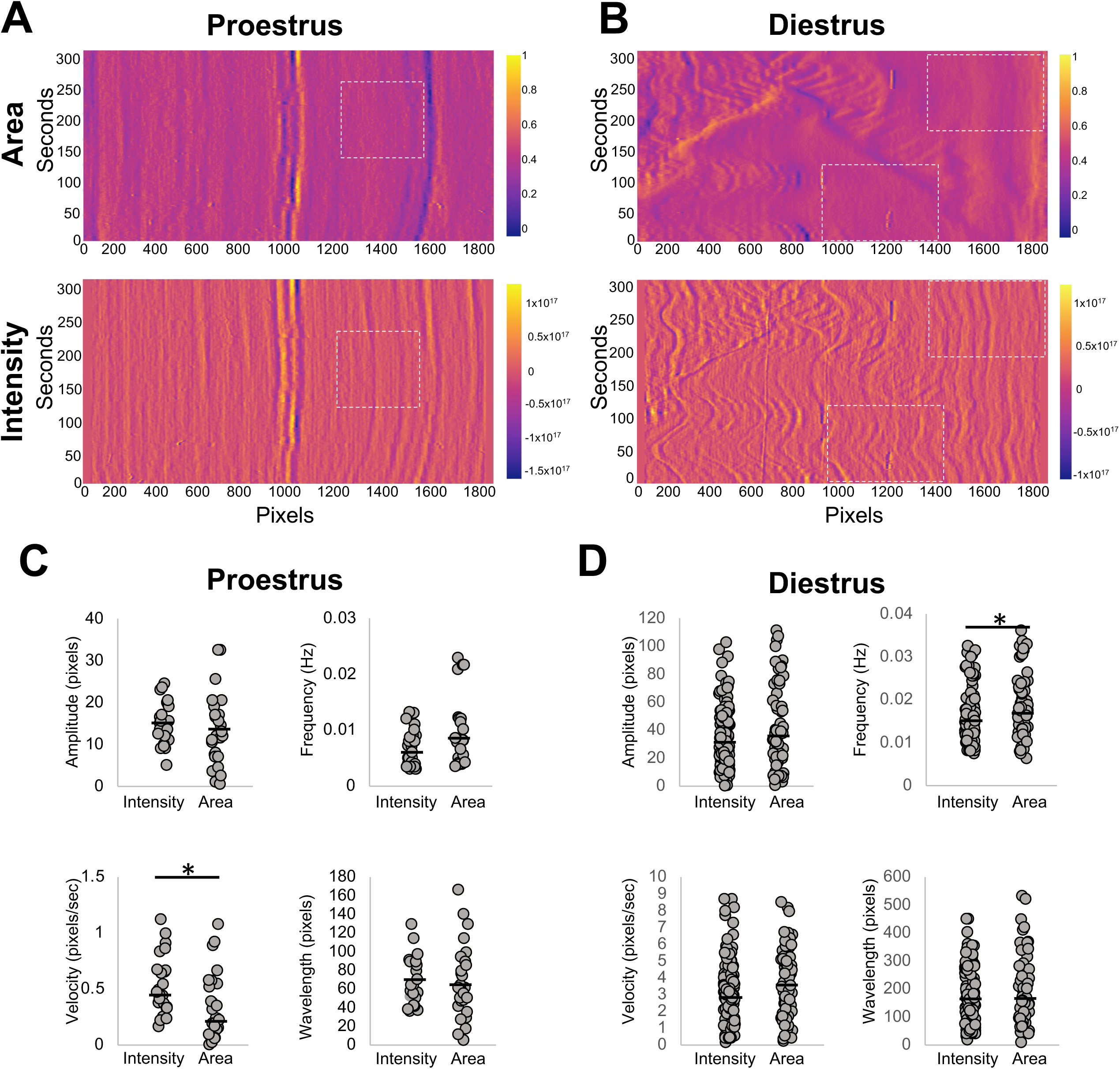
Metrics obtained from 3D area vs 3D intensity plots. Normalized differences in area (top) vs intensity (bottom) in a proestrus (A) and diestrus (B) stage uterine horn. Amplitude, frequency, velocity and wavelength calculations from area and intensity plots from proestrus uterine horn in A (C) and diestrus uterine horn in B (D). White dotted rectangles highlight regions in intensity plots where contractions are observed but corresponding regions in area plots do not show contractions. n= 3 uterine horns for each stage. * P<0.05. Black lines indicate medians.

### Diestrus stage uteri display maximum contractility

To date spatial contractility analysis for the entire uterine horn has only been performed on the stages of estrus cycle in non-pregnant mice. To compare our method to published methods we first applied our method to the different stages of the estrus cycle. We observed that diestrus staged uteri display contractions of the highest frequency (0.015 Hz), and velocity (2.91 pixels/second) compared to each of the other estrus phase (p<0.05 or p<0.0001). Diestrus also showed the highest amplitude (median 29.25 pixels), and wavelength (165 pixels) that was significantly different from proestrus and estrus (p<0.0001) (**Fig. 5A, B, Supplementary Figure 2**). There were similarities between estrus and proestrus stages and diestrus and metestrus stages in trends likely reflecting estrogen dominance of the former and progesterone dominance of the latter (Median values of waveform metrics reported in **Supplementary Figure 2**).

**Figure 5:**
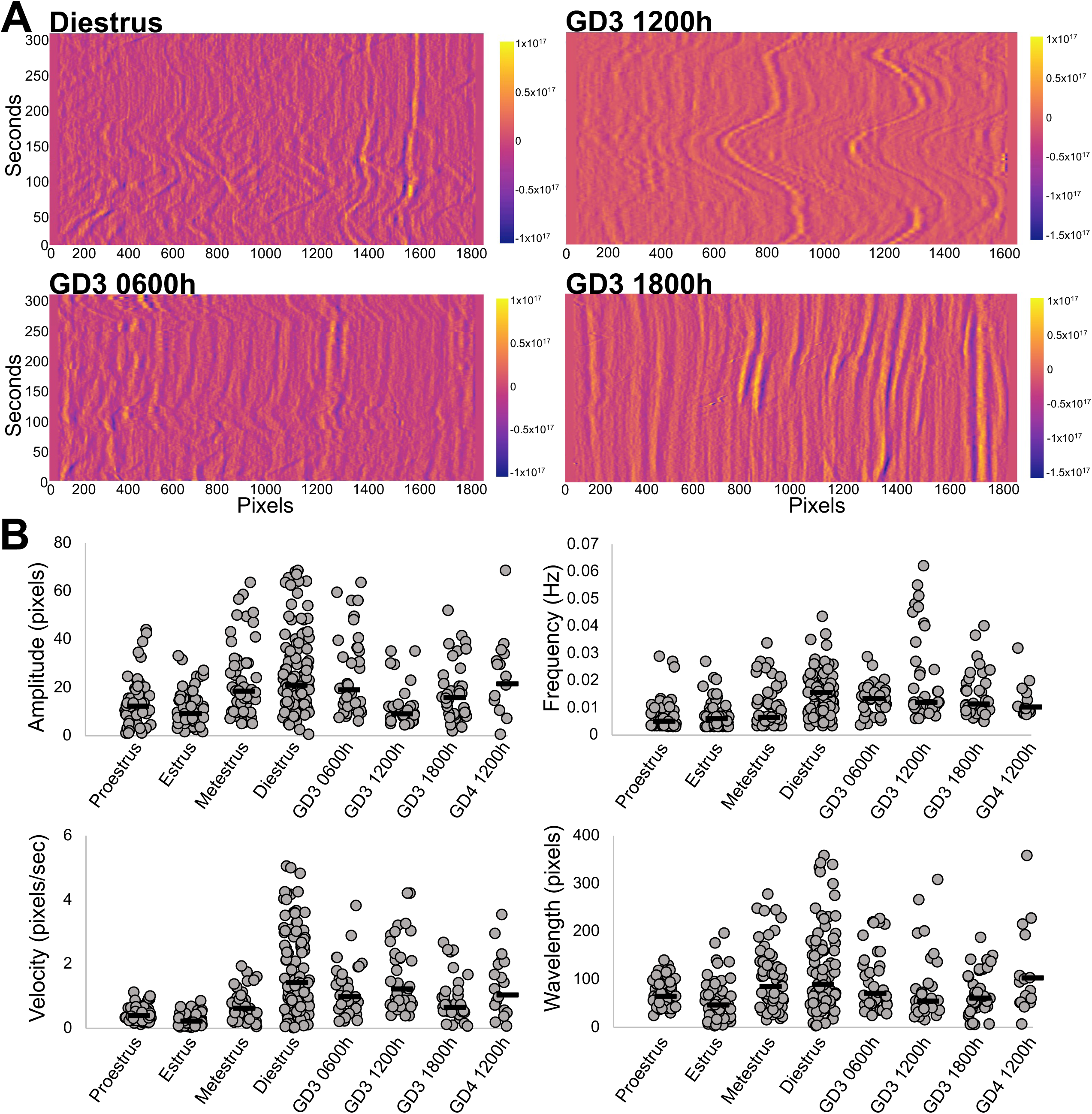
Contractility in pre-implantation stage uteri is closest to the non-pregnant diestrus stage. (A) Example 3D plots for uterine contractility from diestrus, GD3 0600h, 1200h, and 1800h stages. (B) Amplitude, frequency, velocity and wavelength calculations from intensity plots of different uteri. Proestrus (n=3 mice, 6 uterine horns); Estrus (n=3 mice, 6 uterine horns); Metestrus (n=3 mice, 6 uterine horns); Diestrus (n=6 mice, 9 uterine horns); GD3 0600h (n=4 mice, 4 uterine horns); GD3 1200h (n=4 mice, 4 uterine horns); GD3 1800h (n=3 mice, 4 uterine horns); GD4 1200h (n=4 mice, 4 uterine horns). For statistics refer to Supplementary Figure 2 and 3. Black lines indicate medians.

### Pre-implantation pregnancy contractility is most similar to the non-pregnant diestrus stage

The biggest advantage of our method is the ability to assess spatiotemporal uterine contractility in stages of early pregnancy. Pre-implantation, gestational day (GD) 3 of mouse pregnancy displayed contractility most comparable to the diestrus stage of the cycle (**Fig. 5A, B, Supplementary Figure 3**). This is likely due to the progesterone dominance of both GD3 and diestrus stage. GD3 is also the stage where blastocyst staged embryos enter the uterine horn (GD3 0000h), move as clusters to the center of the uterine horn (GD3 0300h - 1200h) and then scatter and space out through the uterine horn (GD3 1200h – 1800h) before they attach on GD4 (0000h) (Flores et al., 2020). We have previously shown that clustered embryo movement is reliant on uterine contractions whereas embryo scattering is independent of uterine contractions. Thus, we assessed contractility corresponding to different phases of embryo movement. We observed that contractions during the early stages of clustered embryo movement (GD3 0600h) display a higher amplitude (median 19 pixels) reflecting active movement of embryos at GD3 0600h. At GD3 1200h the amplitude drops drastically (median 9 pixels) likely because at this time embryos are held in the center of the uterine horn. Intriguingly both GD3 0600h and GD3 1200h display higher velocity of uterine contractions (0.99 and 1.23 px/sec respectively). Velocity of contractions drops during the second phase of embryo movement at GD3 1800h (median 0.66 pixels/second) concurrent with embryo scattering. Similar to diestrus, frequency of uterine contractions stays high throughout pre-implantation embryo movement stages (>0.01Hz) suggesting this wave metric may be responsive to presence of objects inside the uterine horn (eggs in diestrus and embryos in pre-implantation staged uteri). Wavelength on different GD3 time points is comparable to each other but lower than diestrus. Wavelength may be a factor dependent on the diameter of the uterine lumen and may also respond to presence of embryos in the uterine lumen. As the embryos implant and begin to form an implantation chamber at GD4 1200h the waveform metrics are comparable to the diestrus stage and the contractions are more rhythmic throughout the whole horn and spatial variations are not evident.

### Application of contractility method to explain embryo movement patterns in a mouse model with genetic perturbation

Lysophosphatidic Acid (LPA) signals through its receptor LPAR3 in the uterus. This signaling is key for embryo implantation and affects both embryo movement (Flores et al., 2020; Ye et al., 2005) as well as uterine contractility (Hama et al., 2007). However, contractility in LPAR3 mutants was measured using uterine strips and in response to applied tension. Thus, we evaluated spontaneous uterine contractility in WT, *Lpar3^+/-^* and *Lpar3^-/-^* uteri using our method. Intriguingly we observed that when measured globally, metrics for uterine contractility were comparable between WT and *Lpar3^-/-^* uteri and instead the *Lpar3^+/-^*uteri displayed differential contractility from both WT and *Lpar3^-/-^*when evaluating the whole horn (**Fig. 6A, B, Supplemental figure 4**). The contractions present throughout the uterus in *Lpar3^+/-^* uteri at GD3 0600h are of higher velocity (WT=1.34, HET=4.64 and KO=1.11 pixels/second), higher amplitude (WT=19.5, HET=27.5 and KO=17 pixels) and higher wavelength (WT=73; HET=195; KO=71 pixels) compared to both WT and *Lpar3^-/-^* mice. At GD3 1200h *Lpar3^+/-^* uteri display higher frequency (WT=0.0148 ; HET=0.0162 ; KO=0.0141 Hz) but lower amplitude (WT=37.25; HET=20.25; KO=34.5 pixels), lower wavelength ((WT=191; HET=87.82; KO=173.87) and lower velocity (WT=2.96; HET=1.59; KO=2.46 pixels/second) contractions compared to WT and *Lpar3^-/-^* uteri. Our data suggest precocious increase in uterine contractility in *Lpar3^+/-^*uteri compared to both WT and *Lpar3^-/-^* uteri.

**Figure 6:**
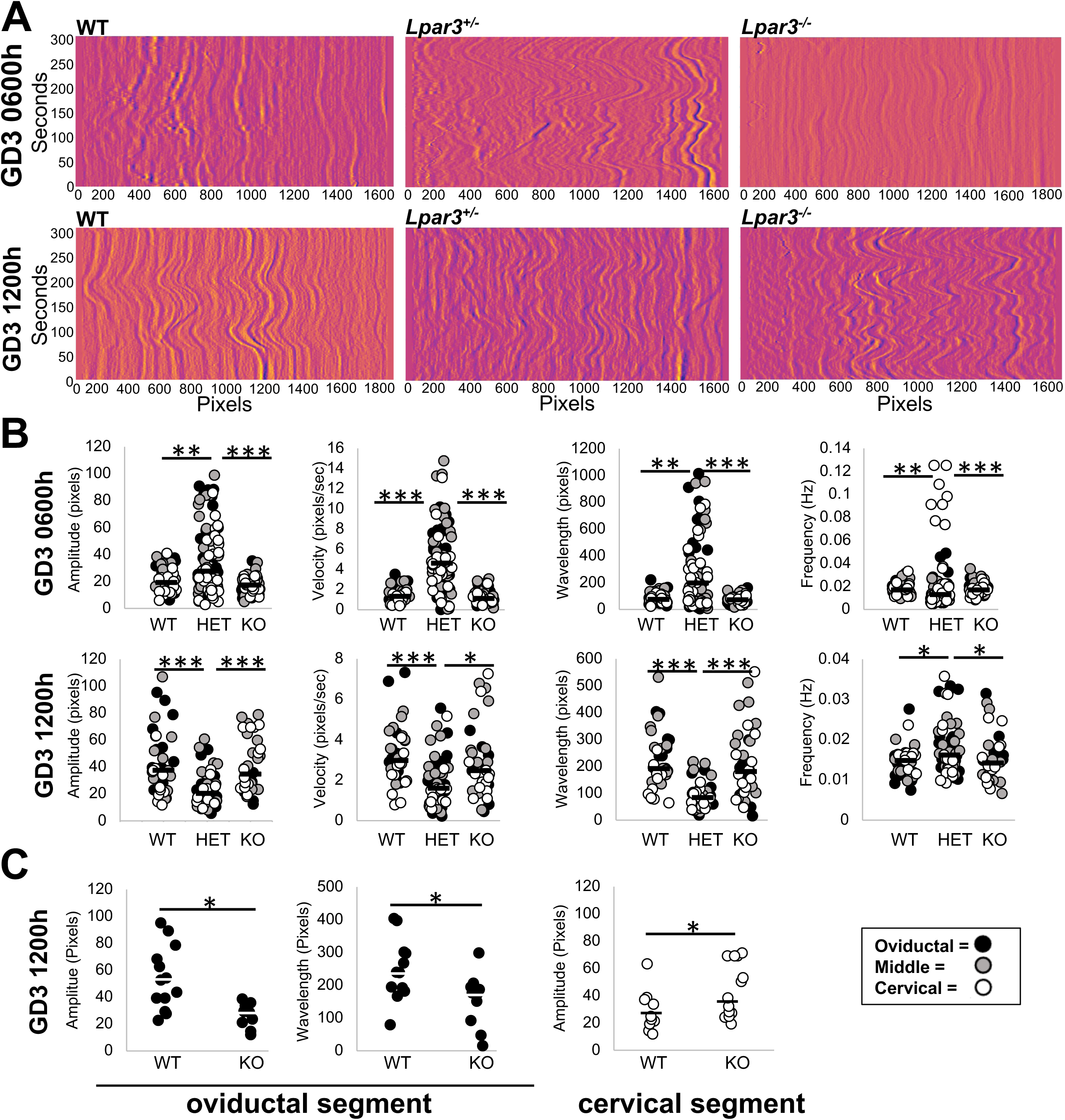
LPAR3-deficient uteri display increased contractility in the preimplantation stages. (A) Example 3D plots for uterine contractility from GD3 0600h and GD3 1200h for wildtype (WT), *Lpar3^+/-^* (HET) and *Lpar3^-/-^* (KO) uteri. (B) Amplitude, frequency, velocity and wavelength calculations from intensity plots of uteri from WT, *Lpar3^+/-^* and *Lpar3^-/-^*uteri. Increased global contractility is observed in *Lpar3^+/-^* uteri compared to WT and *Lpar3^-/-^* uteri. (C) When separated by segment, *Lpar3^-/-^* uteri display decreased contraction amplitude and increased wavelength in the oviductal segment but display increased contraction amplitude in the cervical segment. GD3 0600h WT (n=3 mice, 5 uterine horns); *Lpar3^+/-^* (n=4 mice, 8 uterine horns); *Lpar3^-/-^* (n=3 mice, 6 uterine horns). GD3 1200h WT (n=3 mice, 4 uterine horns); *Lpar3^+/-^* (n=3 mice, 6 uterine horns); *Lpar3^-/-^* (n=3 mice, 4 uterine horns). *P<0.05; ** P<0.001; *** P<0.0001. Black and white lines indicate medians.

While embryo movement patterns for the *Lpar3^-/-^* mice have been studied and it has been shown that embryos in a *Lpar3^-/-^*uteri implant as clusters resulting in implantation loss, embryo movement in the *Lpar3^+/-^* uteri has not been assessed. Since uterine contractions in the first phase of clustered embryo movement are perturbed in *Lpar3^+/-^*mice (**Fig. 6**) we predicted that these mice would display differential patterns of pre-implantation embryo movement. We observed that at GD3 0900h in both controls and *Lpar3^+/-^* mice embryos were approaching the middle of the uterine horn. However, at GD3 1200h, in *Lpar3^+/-^* mice, embryos started to scatter and separate from each other with 48% of embryos in the oviductal segment, 33% in the middle segment, and 19% in the cervical segment while embryos in the control uteri are still clustered in the middle segment with 61% embryos in the middle 31% in the oviductal segment and only 8% in the cervical segment (**Supplemental Figure 5**). Thus, embryos in the *Lpar3^+/-^*uteri show a similar pattern to the controls initially but at mid-day on GD3, their movement appears faster as seen by distribution in three segments.

### Spatial changes in contractility explain embryo movement trajectories in *Lpar3^-/-^*mice

We observed no global differences between WT and *Lpar3^-/-^* mice when evaluating contractility along the whole uterine horn. However, *Lpar3^-/-^* mice display preferential localization of clustered embryos near the cervix at GD3 1200h (Flores et al., 2020; Hama et al., 2007; Ye et al., 2005). Even though global contractility appears to be the same, we determined if contractility in *Lpar3^-/-^* uteri is regionally distinct compared to WT mice. Indeed, we observed differences in contractions between WT and *Lpar3^-/-^*uteri when compared in isolated segments of the uterus (oviductal, middle and cervical segments). At GD3 1200h contraction amplitude is reduced in *Lpar3^-/-^*uteri compared to WTs in the oviductal segment (WT = 52.5, KO =28.75 pixels) and the opposite is true in the cervical segment where contraction amplitude is higher in *Lpar3^-/-^* uteri compared to WTs (WT = 23.5, KO = 38 pixels) (**Fig. 6C, Supplemental figure 4**). At GD3 1200h the wavelength of the contraction in the oviductal segment is also reduced in the *Lpar3^-/-^* uteri compared to the WT (WT= 237.43, KO= 173.87 pixels). These contractility metrics would explain movement of embryo clusters beyond the middle of the uterine horn and closer to the cervix in the *Lpar3^-/-^* uteri compared to controls in the first phase of embryo movement (Flores et al., 2020).

When evaluating just the WT uteri, we noted that at GD3 0600h velocity and wavelength were significantly higher in the middle segment of the uterine horn as compared to the cervical segment, while amplitude and frequency were uniform in all segments (**Supplemental figure 5)**. At GD3 1200h amplitude in the oviductal segment was significantly higher than the cervical segment. At this time, both oviductal and middle segments displayed significantly higher values for velocity and wavelength when compared to the cervical segment while frequency was again similar in all segments (**Supplemental figure 5)**. These data suggest that during contraction dependent embryo movement the cervical end of the uterus is the least active and the middle of the uterine horn is most contractile. Detection of spatial changes in contractility that correlate with observed embryo movement patterns is an example of how our method can be used to learn new facets of uterine contractility during the pre-implantation stages of early pregnancy.

## DISCUSSION

We describe a spatiotemporal method for quantifying uterine contractions. Our method accounts for contributions from both the longitudinal and circular smooth muscle contractions. Most importantly, ours is the first method that allows quantification for both the cycling uterus contractions and contractions in early pregnancy that are key for pre-implantation embryo movement.

### Contractions during estrus cycle

There is conflicting data on which phase of the estrus cycle is most contractile in the rodent. Some report that diestrus is highly contractile and proestrus is the least contractile (Ishikawa and Fuchs, 1978; Wray and Noble, 2008) and some studies show the exact opposite (Griffiths et al., 2006; Houdeau et al., 2003). Crane and Martin, using laparoscopic video recordings suggested that total contractile activity was lowest at pro-estrus, increased at estrus, decreased at metestrus and rose to its highest at diestrus. Using spatiotemporal mapping, Dodds and colleagues (Dodds et al., 2015), clearly indicate that diestrus stage shows maximum contractility. In studying actions of sex hormones on contractions (Wray and Noble, 2008) it is found that estrogen is associated with a reduction in contractions due to reduction in calcium channels that are excitatory for contractile activity and increase in K+ channels that are inhibitory for contractile activity. These authors classify contractile activity as either mechanical or electrical. The electrical activity is low in proestrus, increases in estrus and metestrus and drops in diestrus. However, mechanical activity defined by the propagation of a contraction is highest in diestrus and metestrus. Further, contractile activity can vary by time of day on a certain estrus stage as well (Talo and Kärki, 1976). Alternatively, electrical activity induced contractions in different phases of the mouse estrus cycle suggest that muscle in the estrus stage is most responsive and that in the diestrus stage is least responsive (Houdeau et al., 2003). When force was applied and “spontaneous contractility” under tension transducers was measured (Dodds et al., 2015) contractions in the diestrus phase were quiescent, in the proestrus phase were high frequency phasic, estrus stage were low frequency phasic and metestrus stage were multivariant. In general studies that evoke contractility either using electrical stimulus or using tension transducers conclude that diestrus is not contractile, however when recording spontaneous contractions, we and others (Crane and Martin, 1991b; Dodds et al., 2015) observed maximum contractility during the diestrus phase. Our quantitative data agrees with the imaging-based studies and shows that diestrus stage contractility is highly variable, and the uterus is the most contractile in this stage. We predict that applying external force produces different responses and other oestrus cycle stages are more responsive to force induced contractility compared to diestrus. This suggests that induced contractions may indicate ability of the muscle to respond to stimuli and this may be different from the native spontaneous contractions.

#### Frequency

When comparing numerical values of different waveform metrics, Dodds and colleagues (Dodds et al., 2021) reported frequency at 0cm water distension to be 1.2 contractions per minute at proestrus which is comparable to data from Zhang and colleagues (Zhang et al., 2019) that described an overall dominant frequency of ∼1 per minute from in vivo recordings. However, the in vivo study also shows that there is a range of frequency from 0.008Hz – 0.029Hz. Our proestrus stage data shows contractility with frequency of 0.003 to 0.029Hz that aligns well with the range of frequency detected from in vivo studies.

#### Velocity

Dodds and colleagues in their first report, suggest contraction velocities from spatiotemporal mapping to be (Dodds et al., 2015) a mean value of 1.3mm per second (proestrus); 0.9mm per second (estrus); 1.2mm per second (estrus) and 0.7mm per second (diestrus). These values are significantly lower than those reported by Dodds and colleagues in 2021 in the proestrus stage reported as 7.4mm per second (Dodds et al., 2021). Our data suggests a maximum median velocity of ∼2.9 pixels per second which translates to 0.026mm per second in the diestrus stage. Thus velocity comparisons do not match up across studies. Differences in waveform metrics could be due to the use of different buffer or mouse background. Our study uses PBS and mixed background mice while Dodds and colleagues use Krebs buffer and C57Bl6 mice. Thus, comparisons across genotypes or time points with mice on the same background may be more useful and accurate rather than comparing across studies.

#### Amplitude

Maximal strength of the contractile wave of amplitude has been measured using different methods including applying mechanical tension (Dodds et al., 2015), based on pressure recordings (Dodds et al., 2021) or a single maximal amplitude throughout the uterine horn was calculated using Fast Fourier Transformation (FFT) (Zhang et al., 2019). We cannot compare our spontaneous contractility data to tension induced contractility. FFT, can only be applied if a single wave propagates throughout the entire uterine horn. Our goal was to calculate local spatial wave metrics to discern contributions of muscle contractility to embryo movement in the pre-implantation phase and we did not calculate a maximal amplitude as has been reported in prior methods.

#### Regional differences in contractility

Some studies have reported that caudal uterus (closer to cervix) is more contractile than the rostral uterus (closer to the oviduct) (Houdeau et al., 2003). Stage specific spatial contractility has also been described where in diestrus, contractility is highest in oviductal segment while contractility in proestrus is highest in the cervical segment (Ishikawa and Fuchs, 1978). Dodds and colleagues (Dodds et al., 2015) report that contractions usually originate from the oviductal region in proestrus and estrus and have multiple origin points throughout the horn in metestrus and diestrus. Our 3D area and intensity plots did not support preferential contractile activity in the oviductal or cervical end during proestrus.

We observed that contractility is highest in progesterone dominant diestrus phase and in the pre-implantation stages. This idea is supported by muscle specific progesterone receptor (PGR) knockout that causes embryo retention in the oviduct and impaired myometrial adaptation to pregnancy (Wu et al., 2022). PGR-deficient uteri show reduced response to oxytocin induced contractility and show reduced expression of calcium homeostasis genes. Transcriptomic analysis suggests that major matrix and muscle genes are regulated by PGR signaling including Myocd and Ccn2. However, to prevent contractions during pregnancy, evolutionarily high progesterone signaling must be linked to uterine quiescence. This apparent discrepancy can be explained by the fact that PGR has two isoforms PGRA and PGRB. While PGRB promotes relaxation, PGRA promotes contractility in human uterine smooth muscle (Peavey et al., 2021). Overexpression of PGRB increases gestational length and overexpression of PGRA increases uterine contractility without affecting gestational length. Transcriptional analysis suggests that PGRB induces muscle relaxation and PGRA is proinflammatory. Thus, depending on which isoform of PGR is expressed and in what tissue, progesterone may induce differential contractile responses as needed for cyclicity and pregnancy success.

### Comparison to other methods for contraction measurement in pregnant state

While there has been a lot of investigation of uterine contractility during different stages of the estrus cycle and during late-term pregnancy, information on contractility during the peri-implantation phase is sparse. Oviductal myogenic contractions have been measured and shown to be responsible for egg/embryo movement during early pregnancy (Dixon et al., 2009). Using spatiotemporal mapping oviductal spontaneous contractions were measured although the authors showed that additional contractility can be induced by applying mechanical force. They were simultaneously able to trace the edge of the oocyte and determine that oviductal spontaneous peristaltic like contractions drive movement of the egg. These myogenic contractions and resulting egg movement were inhibited by calcium blockers and were not perturbed when blocking nerve activity. Further while blocking contractility blocks egg and embryo movement, small particle movement (<25μm) that relies on ciliary activity was not blocked (Ward et al., 2022). Oviductal myogenic contractility also relies on Interstitial Cells of Cajal (ICC). Blocking ICC function by Kit antibody, or perturbation of ICC networks by bacterial infection, both block propagating contractions. There is evidence of ICC in the uterus (Dodds et al., 2015), however whether ICC provide pacemakers and initiation of contractility has not been proven.

To conclusively prove that uterine peristaltic spontaneous contractions directly cause uterine embryo movement, simultaneous recordings of embryo movement and uterine wall would be necessary. However, the uterine smooth muscle is ∼500μm thick preventing simultaneous live imaging of the muscle activity and the embryo movement. Thus, in the current study we are using our previously published patterns for embryo movement (Flores et al., 2020) and contractility recordings (this study) to evaluate contributions of contractions to embryo movement over time. Our data suggests that early on GD3 of mouse pregnancy, during the first phase of embryo movement, contractile waves are not uniform throughout the horn. With spatial analysis we determined that the middle of the uterine horn is most contractile while the cervical end of the uterus showed least contractile activity. As, the embryos enter the second phase of movement where they scatter and space out, contractions along the entire horn become more rhythmic and uniform. These data would support a more active role for uterine contractions during the first phase of movement as has been observed by use of muscle relaxants in vivo (Flores et al., 2020).

### Comparison of image analysis methods

Prior studies that measure contractions based on video-recording generate kymographs or spatiotemporal maps that were initially developed for the intestine (Hennig et al., 1999) and then applied to the oviduct (Dixon et al., 2009), and the non-pregnant uterus (Dodds et al., 2015; Dodds et al., 2021). These methodologies produce very similar data to our area based 3D plots. Zhang and colleagues (Zhang et al., 2019) used 8 different equally spaced locations along the uterine horn to assess the movement of the uterine horn edges away from a central line through the uterine horn and then applied an area based calculation. Our method utilizes changes along the length of the uterine horn and accounts for changes in area but we also evaluate changes in contractility along the uterine horn using tomato intensity. These changes in intensity are particularly useful to study spatial distribution of contractions along the uterus in pre-implantation stages of pregnancy.

Below we summarize the advantages and limitations of our method:

#### Advantages

Our method presents the following advantages over previously published methods:

(i) We can record any length of uterine horn and thus we are able to record contractions from both CD1 (**Fig. 3, 4**) and C57Bl6 mouse background (**Fig. 6**). The limit for uterine horn length is determined by the objective lens used for capturing the tomato+ images.
(ii) We are able to image diestrus and early pregnancy stages that have stronger and more variable contractions.
(iii) Using intensity plots for measuring contractions, we have the spatial resolution to separate effects on different segments of the uterus – closer to the oviduct, middle of the uterine horn and closer to the cervix.
(iv) Since our recordings are ex vivo and not in vivo, in our datasets contractility is not influenced by an anesthetic.
(v) Despite being an ex vivo method our initial measurements for frequency of contractions match up with the in vivo data (Zhang 2019).
(vi) Our method is also amenable to short-term manipulation in the culture dish with fast acting small molecule inhibitors and activators. Alternatively, mice can be treated in vivo for longer acting molecules such as ovarian hormones and still be assessed later.

#### Limitations

Although our method presents an advance and allows recording of pre-implantation stage uteri there is room for improvement of the method. Following limitations are noted:

(i) Although the area plots can be obtained from any mouse line, for getting the intensity data tomato allele needs to be bred with the mice under investigation.
(ii) Animals need to be sacrificed thus the same animal cannot be imaged repeatedly for different stages.
(iii) Pinning the uterine horn in its in vivo configuration is a manual component of the method. Stretching the uterus too much will prevent in vivo like measurements and leaving slack in the uterus may prevent the computational workflow to be applied to the captured images.
(iv) Our recordings are done promptly after the animal is sacrificed and we record contractility for five minutes. Any waves that have a frequency larger than five minutes will be missed.
(v) The buffer we image in can impact contractility and this should be kept in mind when comparing data generated from our method to results from other studies.
(vi) Our method allows for relative quantification of two stages and absolute values of waveform metrics should be used cautiously. Our method was used to measure contractions across different mouse backgrounds and we did observe that at pregnancy stage GD3 1200h, contractility metrics for CD1 mice (**Fig 5**) differed from wildtype litter mates on C57Bl6 background (**Fig. 6**). These data suggest variations in contractility due to genetic backgrounds and further emphasize that these methods should be used to compare tissues on the same genetic background.
(vii) Although we obtain 3D intensity plots and a lot more data, we are still limited by manual calling of waves and calculating metrics based on those calls. Further, lots of smaller waves can shadow the larger waves in the 3D plots making it harder to analyze larger waves in a highly contractile uterus. For both these reasons the method of calling waves needs to be automated and is a subject of future investigations.

### LPAR3: a stimulant or relaxant for uterine contractions?

LPA-LPAR3 signaling pathway has been implicated in embryo movement and uterine contractility (Flores et al., 2020; Hama et al., 2007; Ye et al., 2005). *Lpar3^-/-^* uteri display embryo clusters that move past the middle of the uterine horn in the first phase of movement and a failure of embryo clusters to separate in the second phase of embryo movement. In vitro mechanical transducer-induced assays suggested that while control uteri can respond to LPA-LPAR3 agonist, *Lpar3^-/-^*uteri fail to respond to the agonist as measured by contractility however these uteri contract in the presence of acetyl choline (Hama et al., 2007). Further, LPA can dose dependently increase contraction amplitude in estrus stage rat uteri in in vitro uterine strips and in vivo (Nagashima et al., 2023; Tokumura et al., 1980). Additionally, a study with gilts (pig uteri) suggested that LPA signaling enhances contractility in early pregnancy uteri but not non-pregnant uteri (Markiewicz et al., 2012). All of these studies suggest that LPA-LPAR3 pathway activation is associated with increase in contractions. However, movement of embryo clusters past the middle of the uterine horn and closer to the cervix in the contraction dependent phase of embryo movement would predict that spontaneous contractions are increased in LPAR3-deficient uteri. Our method demonstrates a spatial disruption in *Lpar3^-/-^*uterine contractions. While in wildtype uteri, contractions are active in the oviductal segment of the uterine horn, contractions in *Lpar3^-/-^* uteri are higher in the cervical segment of the uterine horn. Further, *Lpar3^+/-^* uteri also display higher contractility that correlates with faster movement of embryos in these uteri. It is critical to note that while the increase in contractility was observed at GD3 0600h (**Fig. 6**), the faster embryo movements were observed a little later at GD3 1200h (**Supplementary Figure 5**). These data suggest that there are other factors in addition to contractility that may fine tune embryo movement patterns. Thus, loss of LPA-LPAR3 activity may be related to enhanced contractility in a regional manner in the first phase of embryo movement. LPA agonists and antagonists can affect contractility differently in pregnant uteri when compared to cycling uteri (Markiewicz et al., 2012), thus there may be a hormonal component in regulating when LPA signaling activates contractions or inhibits contractions.

### Conclusions and Future Directions

Contractility, especially in the context of object movement, is complex and has many facets, including what kind of stimuli is used to induce contractility, what the hormonal milieu is, and the least studied in the context of uterine contractions is whether the diameter of the uterine lumen impacts propagation of uterine contractions. Proestrus and estrus staged uteri are fluid filled due to estrogen regulated luminal epithelium secretions (Kalyne and Bruce, 2014). On the other hand, diestrus staged uteri show narrower luminal openings (Kalyne and Bruce, 2014) which likely support contractile waves to travel along greater distances. Thus, hormonal impacts cannot be considered independent of the effects of ovarian hormones on the luminal epithelium structure and volume and nature of luminal epithelial secretions. Theoretically, estrogen in proestrus and estrus stages may show reduced propagation of contractile waves due to an open lumen, however supplementing with estrogen during progesterone-dominant pre-implantation may increase contractility due to a closed lumen. This idea is supported by our observation that embryos travel farther along the uterine horn when treated with estrogen during pre-implantation stages(Lufkin et al., 2023). Progesterone may permit luminal closure but by itself may have an inhibitory effect on how far an individual contraction travels. Thus, several open questions remain. For example how do absolute levels of ovarian hormones or an estrogen:progesterone ratio regulate uterine lumen structure, fluid accumulation in the lumen and movement of objects through the lumen? Do objects such as eggs and embryos in the uterine horn act as pacemakers to trigger spatially distinct contractile waves? Our method can now allow researchers to start addressing some of these scenarios. In the long-term, understanding how contractility regulates early embryo movement and how factors such as ovarian hormones or signaling molecules such as LPA regulate uterine contractions can be useful in modulating contractility during embryo transfer to improve outcomes for invitro fertilization and artificial reproductive technologies in the clinic.

## CONFLICT OF INTERESTS

The authors declare no conflict of interests.

## AUTHOR CONTRIBUTIONS

M.D, D.F. and R.A. designed the experiments; M.D, D.F., and L.Z. performed experiments; O.Z. and D.F., wrote the code for data analysis, M.D, L.Z., S.A., R.M., O.Z., and R.A. analyzed the data. M.D., L.Z., and R.A. interpreted the results; M.D., L.Z., and R.A. wrote the manuscript.

## FUNDING

We acknowledge support from grant NIH R01HD109152 to RA and NICHD T32HD087166 to LZ.

## ACKNOWLEDGEMENTS

We are also grateful to Drs. Asgerally Fazleabas, Nataki Douglas and Gregory Burns for critical analysis and research discussions.

## DATA AVAILABILITY STATEMENT

Data generated in the manuscript is available upon reasonable request to the corresponding author.

---

**Supplementary Figure 1:**
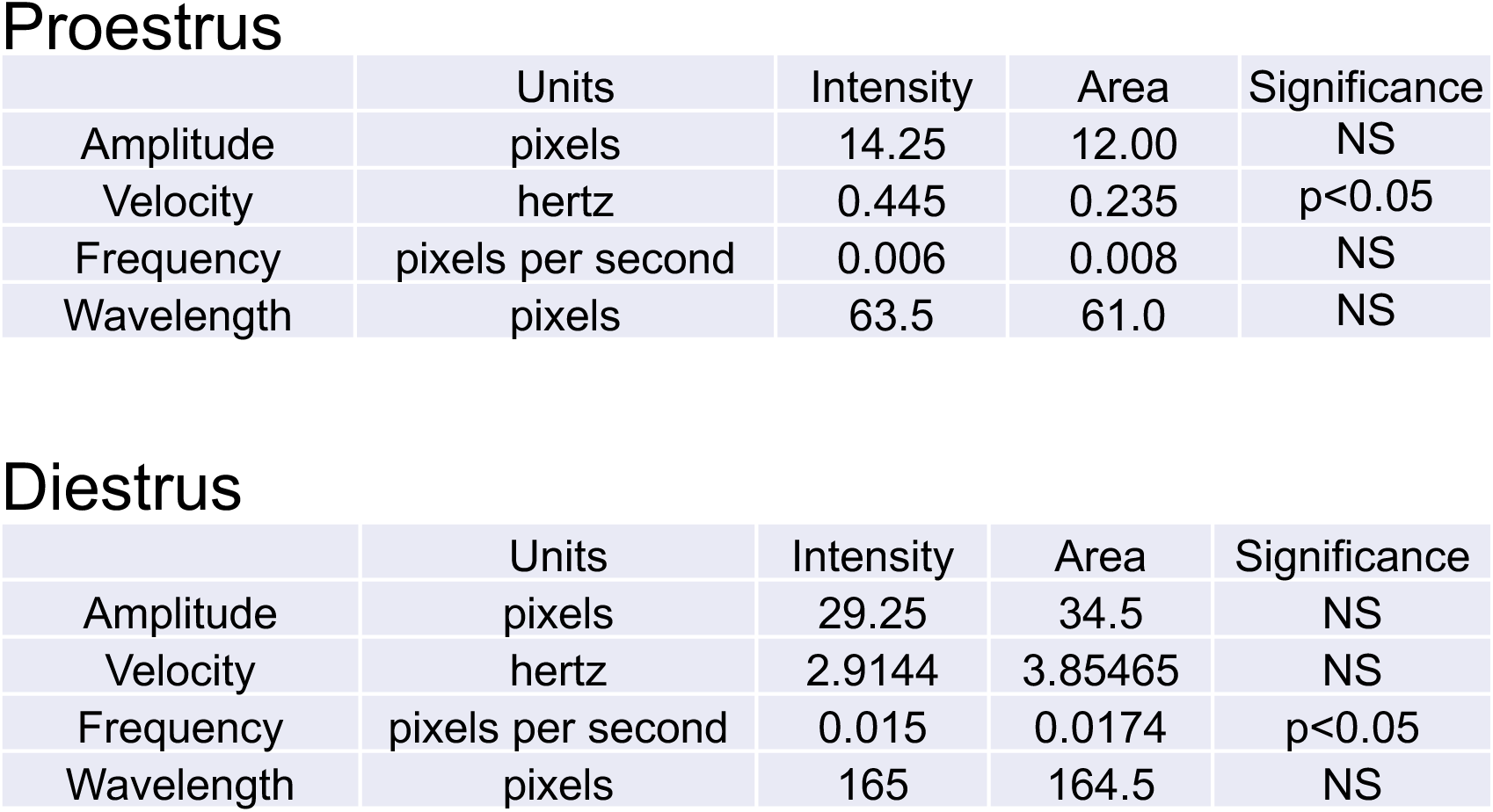
Median values of waveform metrics from 3D intensity plots or 3D area plots and statistical differences between calculated metrics. (Stats for Figure 4)

**Supplementary Figure 2:**
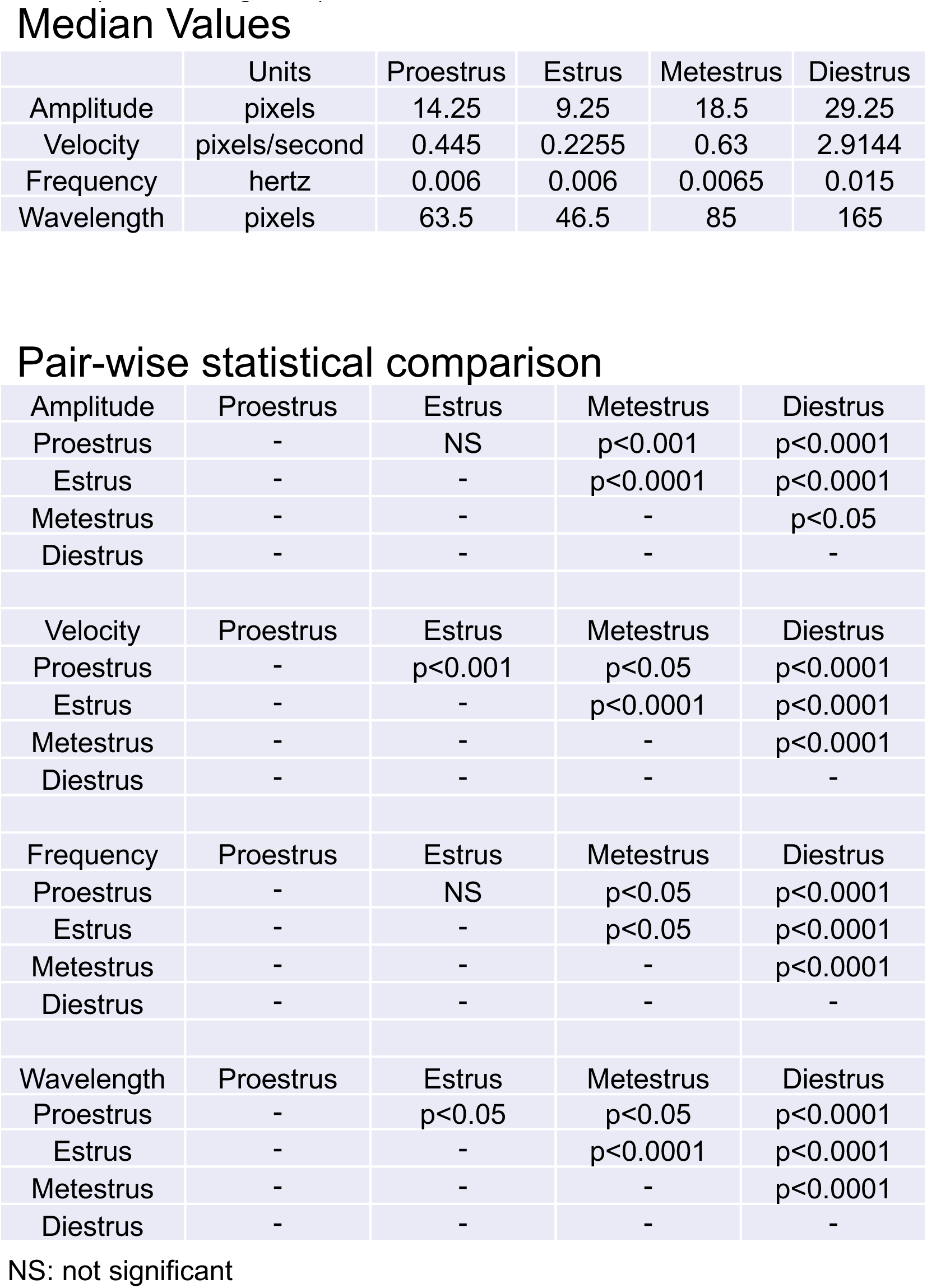
Median values of waveform metrics from 3D intensity plots for different phases of estrus cycles followed by pairwise comparisons for statistical differences between calculated metrics. (Stats for Figure 5)

**Supplementary Figure 3:**
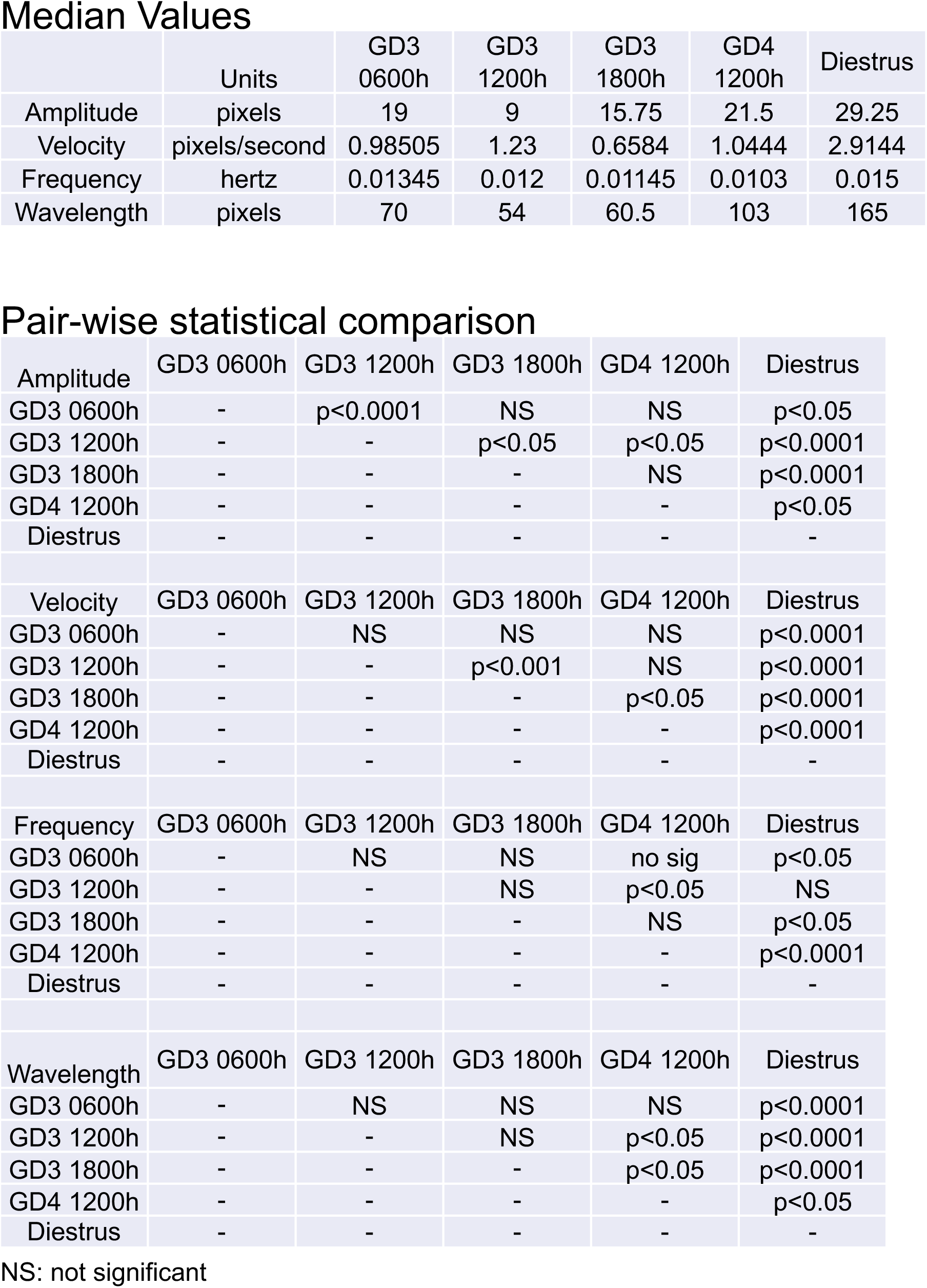
Median values of waveform metrics from 3D intensity plots for pre-implantation pregnancy time points followed by pairwise comparisons for statistical differences between calculated metrics. (Stats for Figure 5)

**Supplementary Figure 4:**
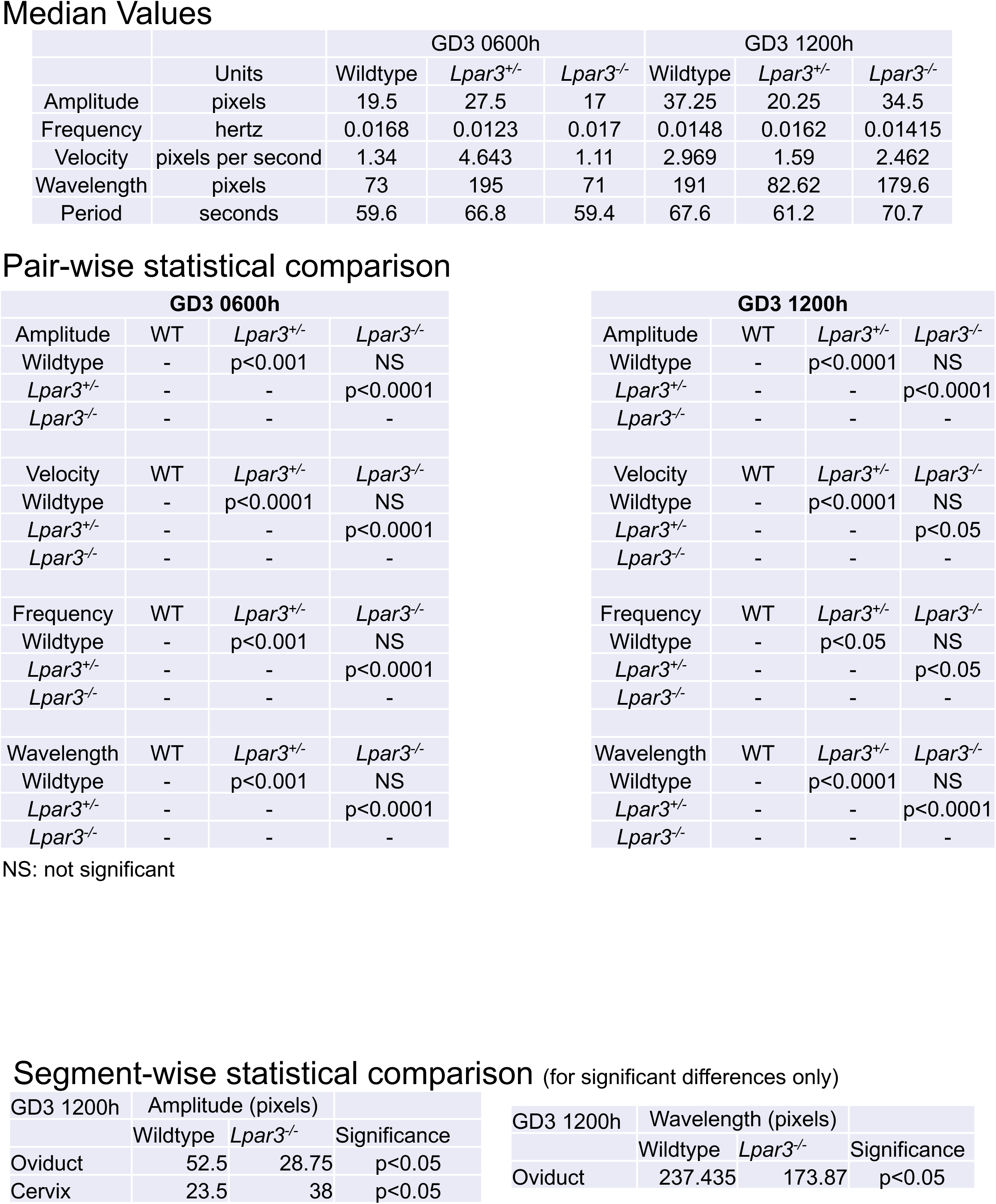
Median values of waveform metrics from 3D intensity plots for pre-implantation pregnancy time points in wildtype, *Lpar3^+/-^* and *Lpar3^-/-^* uteri followed by pairwise comparisons for statistical differences between calculated metrics. (Stats for Figure 6)

**Supplementary Figure 5:**
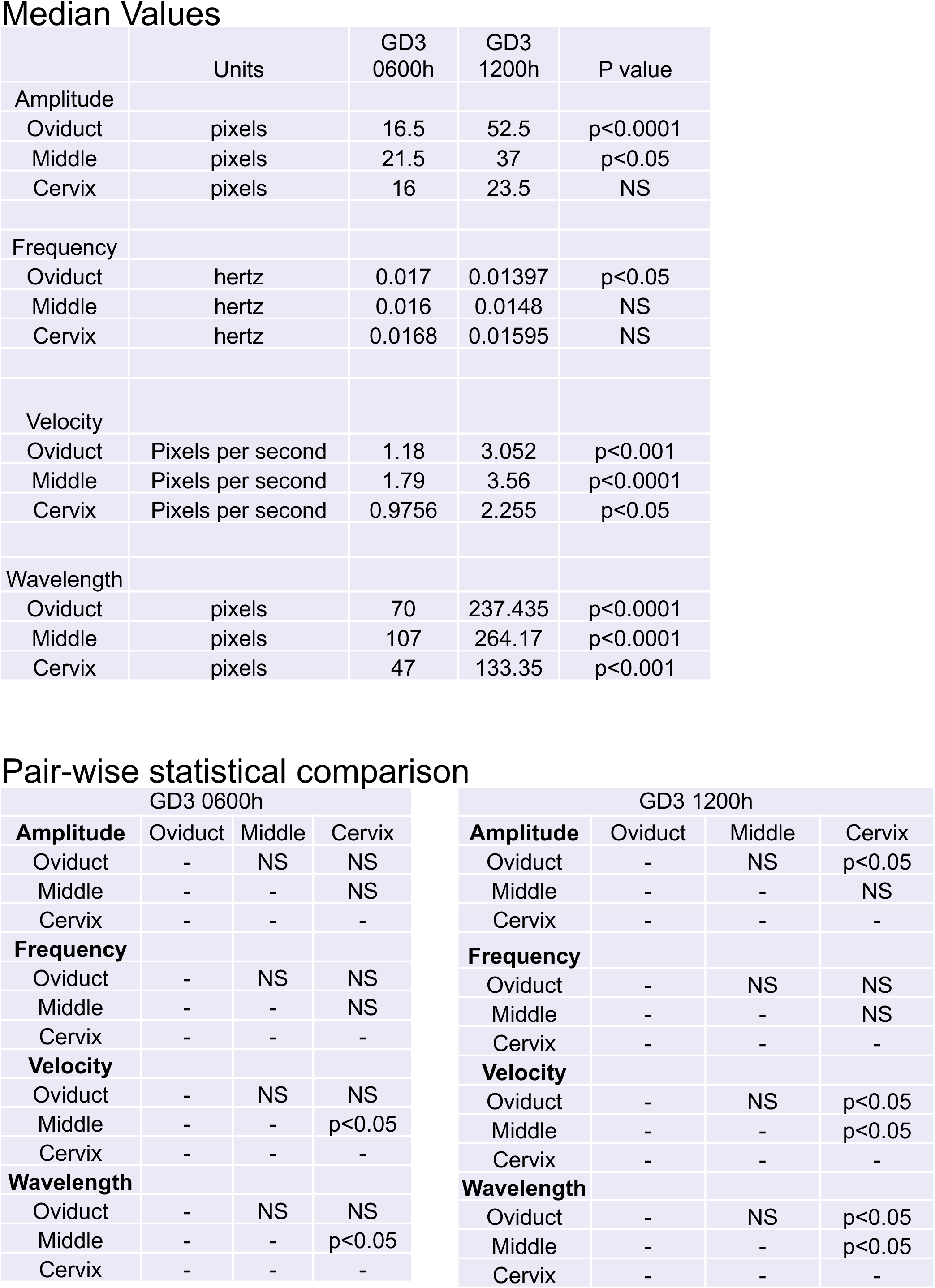
Median values of waveform metrics from 3D intensity plots for pre-implantation pregnancy time points in wildtype but split in different segments of the uterine horn.

**Supplementary Figure 6:**
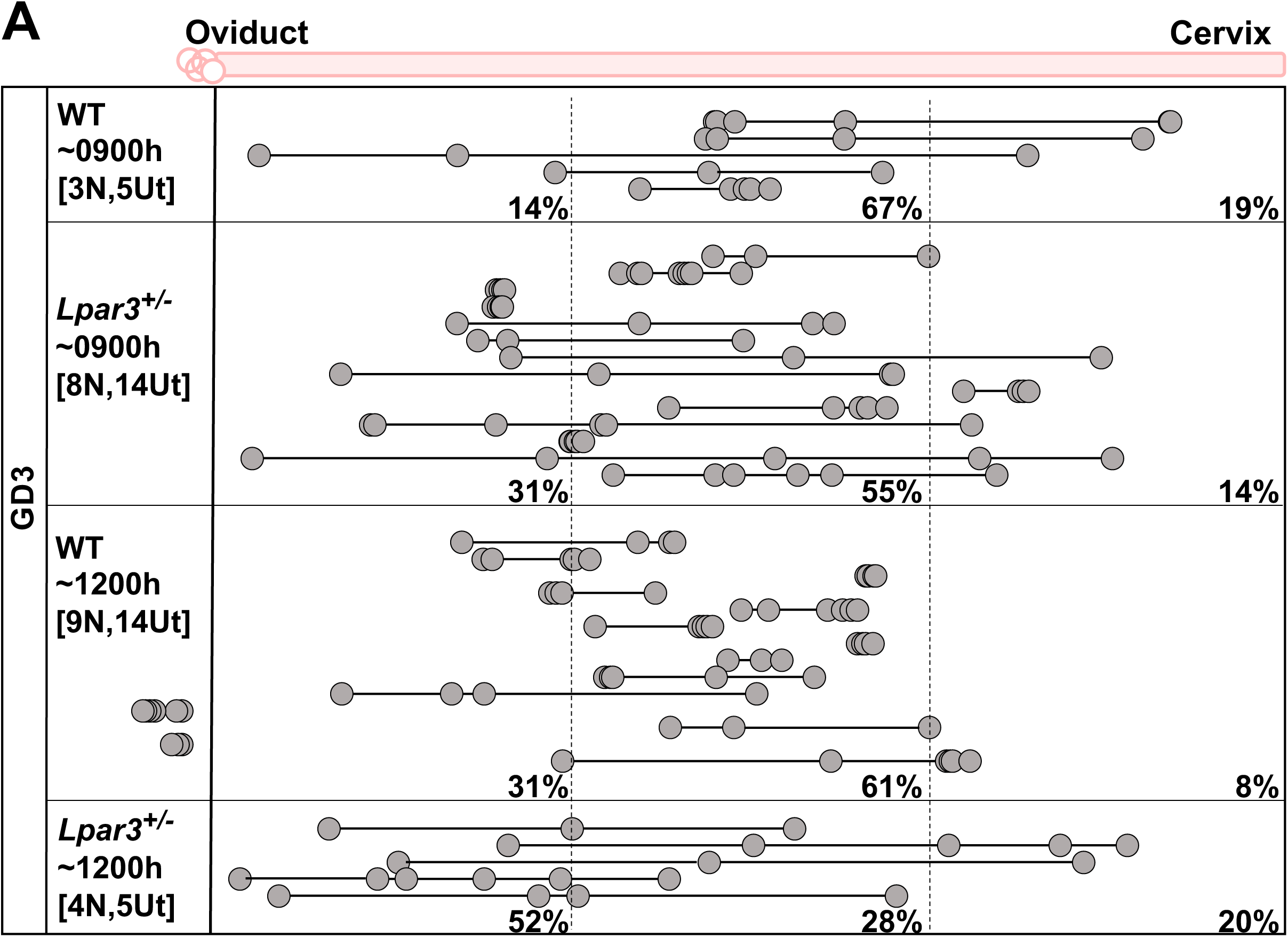
Embryo location suggests faster movement of embryos in *Lpar3^+/-^* uteri. Embryos in WT uteri display embryo clusters in the middle of the uterine horn at GD3 0900h at GD3 1200h. *Lpar3^+/-^* uteri display clustered embryos at GD3 0900h, however at GD3 1200h, embryo clusters are already separated in the three uterine horn segments suggesting faster embryo movement likely due to increased contractility in these uteri.

## Notes

### Competing Interest Statement

The authors have declared no competing interest.

