## Supplementary Methods for "Imaging the dynamics of uterine contractions in early pregnancy"

----------------------------------------- Summary of the method with images --------------

STEPS

<https://docs.opencv.org/4.x/d4/d86/group__imgproc__filter.html#gaabe8c836e97159a9193fb0b11ac52cf1>

-- Smooth with Gaussian in time

cv2.GaussianBlur ( COLUMN, 3, 3)

-- Smooth with Gaussian in Space

cv2.GaussianBlur ( COLUMN, 5, 5)

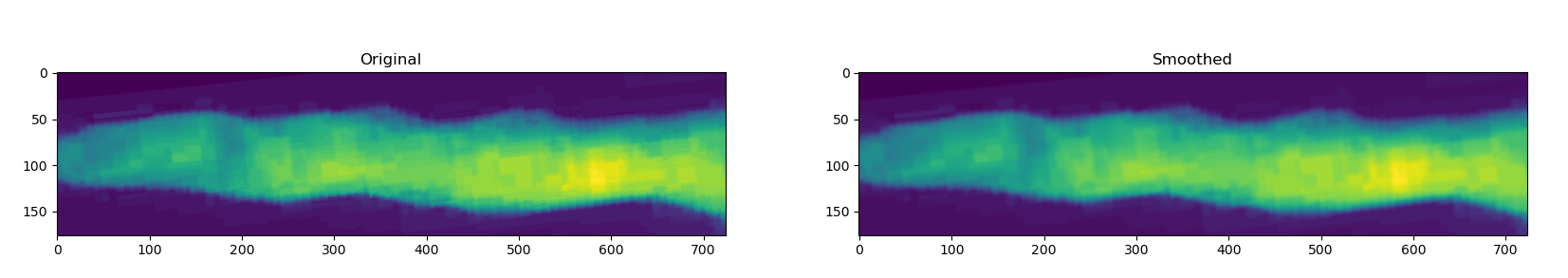

--- Compute Sobel from Smooth video

<https://docs.opencv.org/3.4/d2/d2c/tutorial_sobel_derivatives.html>

Sobelx = cv2.Sobel( img, 0, 1, kernel_size = 5)

Soboely = cv2.Sobel( img, 1, 0, kernel_size = 5)

Sobel Output = 2*sobelx + sobely

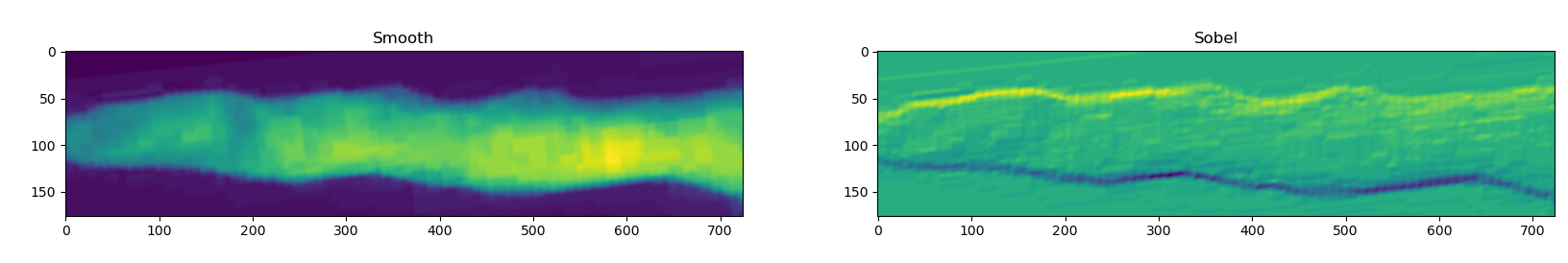

-- Detect top position from the cumulative sum of the Sobel and a predefined threshold. Manually tuned for each video with a default value of **1600**

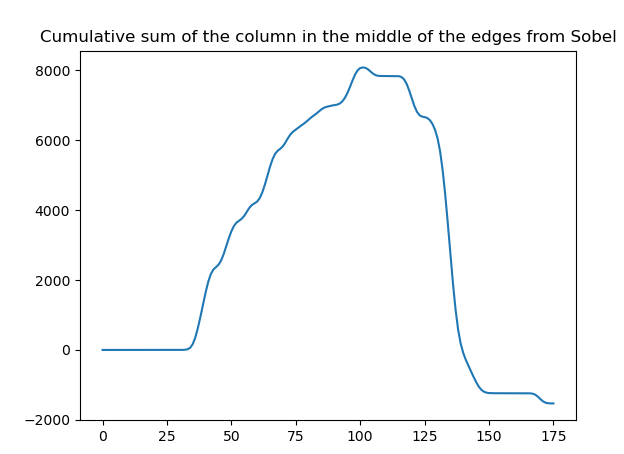

-- Detect bottom position from the ‘flipped’ cumulative distribution, manually adjusted with a default threshold of 1900.

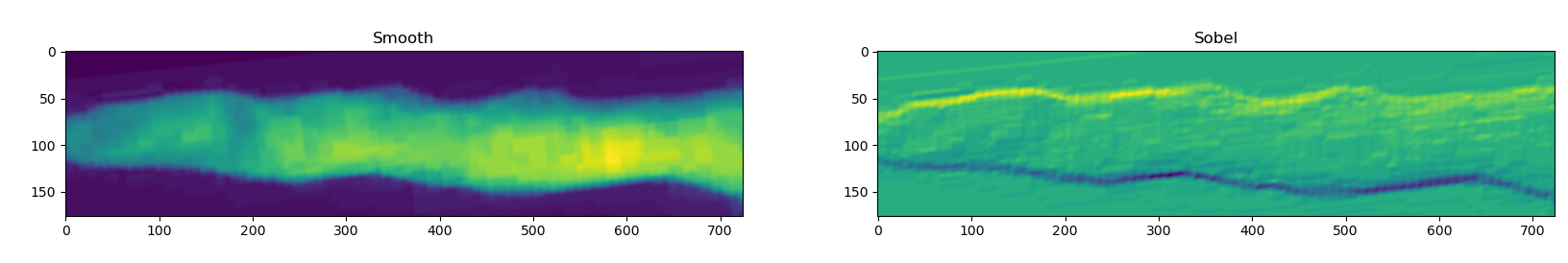

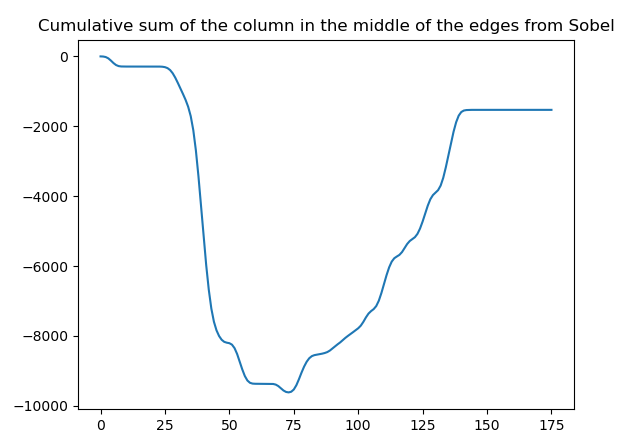

Output

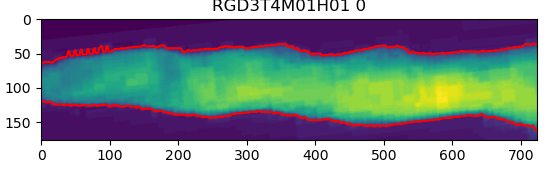

-- Apply median filter to the locations. With default kernel size of 5

scipy.signal

medfilt(top_pos[cur_frame], median_filt_size_top)

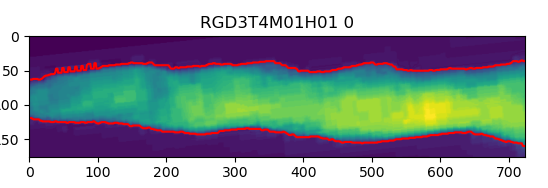

-- CubicSplines

Interpolates the curves using cubic interpolation. The number of points is around **cols/1000 * 50** for the top and **cols/1000*100** at the bottom. Around something like 36 and 70 in the example below.

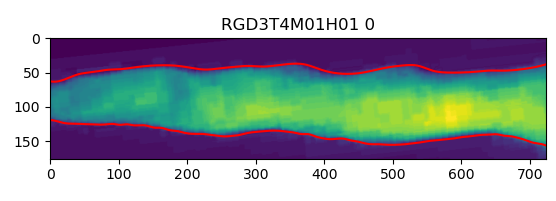

-- Saves mean intensities, bottom, top, and area (bottom - pos)

======================== Second file ==========================

For each previous sequence (area, top, bottom, etc) computes:

1. (name should be space variations) Vertical edges **(sobel 0,1 ksize=21)**→ This is used to detect space variations (by column) independent of the areas ‘far away’ from the current location.
2. (name should be time variations) Horizontal edges (**sobel 1,0 ksize =31)** → This is used to detect changes between time (not very clear most of the time)
3. Filtered image
   a. Compute low frequencies from a running mean of 40 columns
   b. Removes the low frequencies from the original
   c. Smooths a curve with a running mean of 3 columns

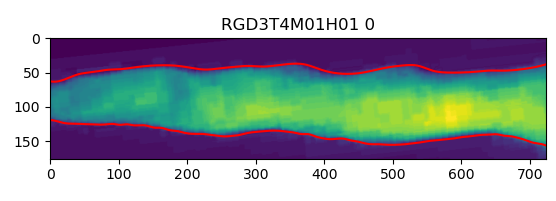

Area

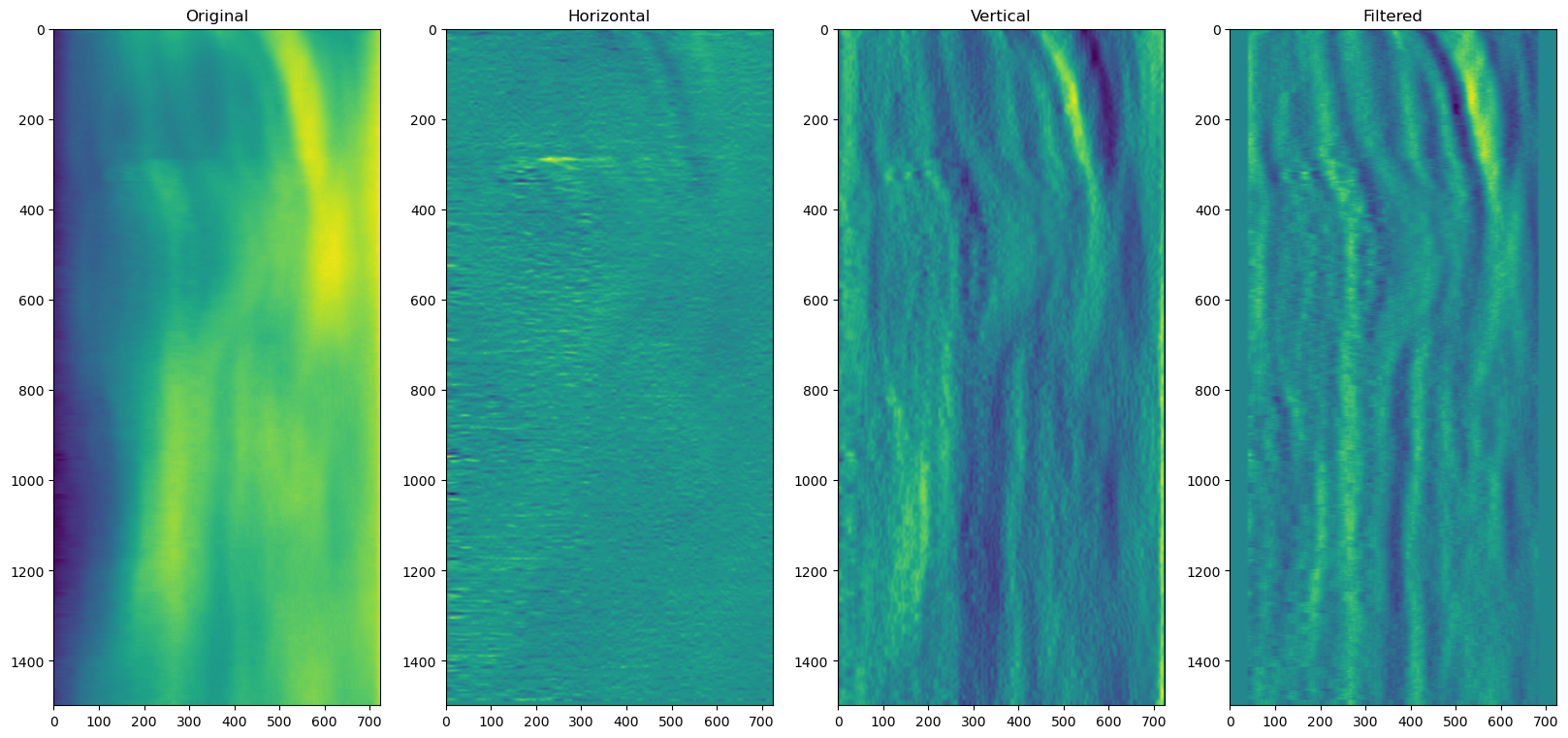

Intensity → Number of cells

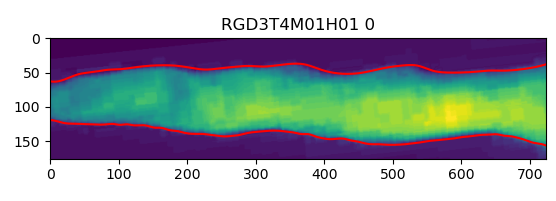

Mean intensities

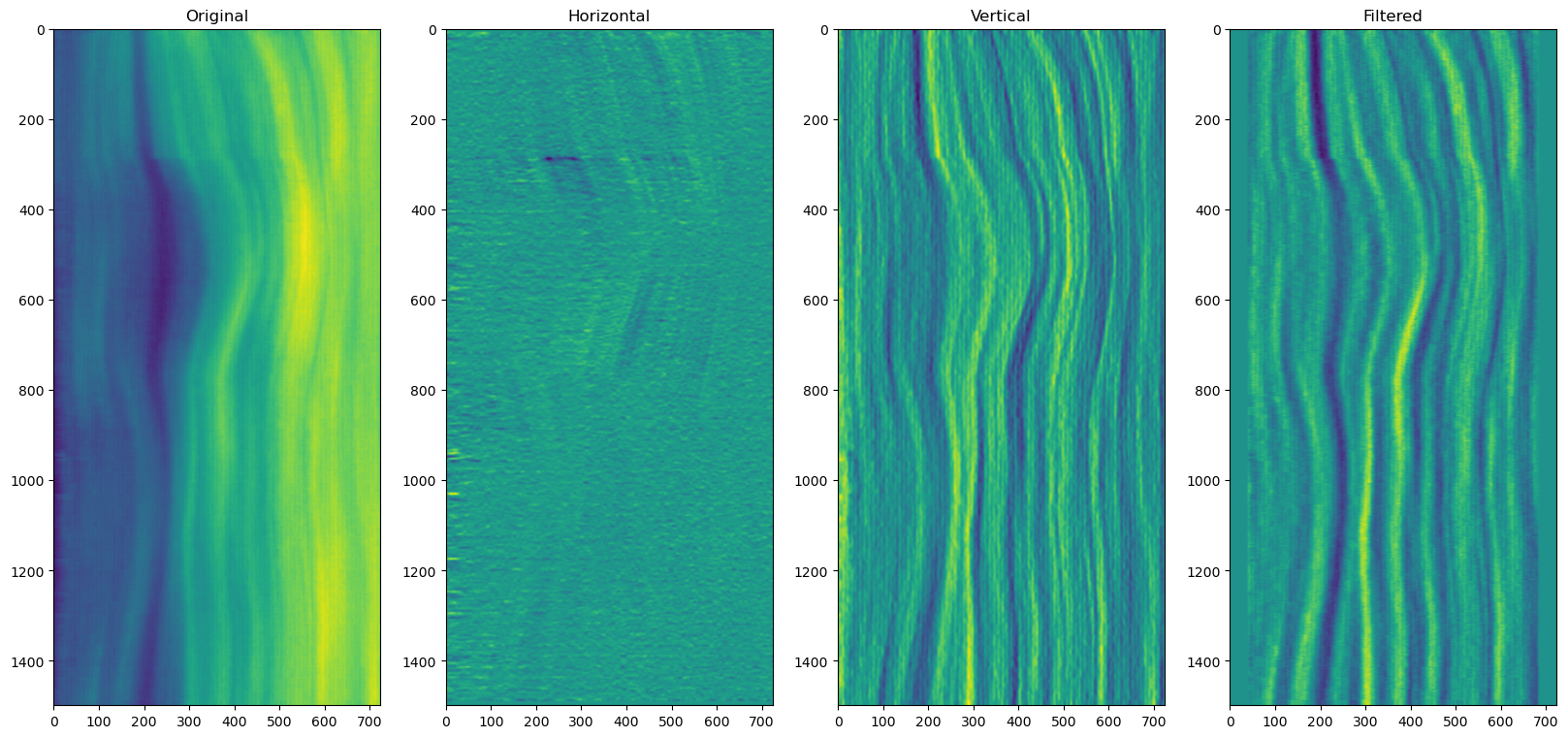

Bottom

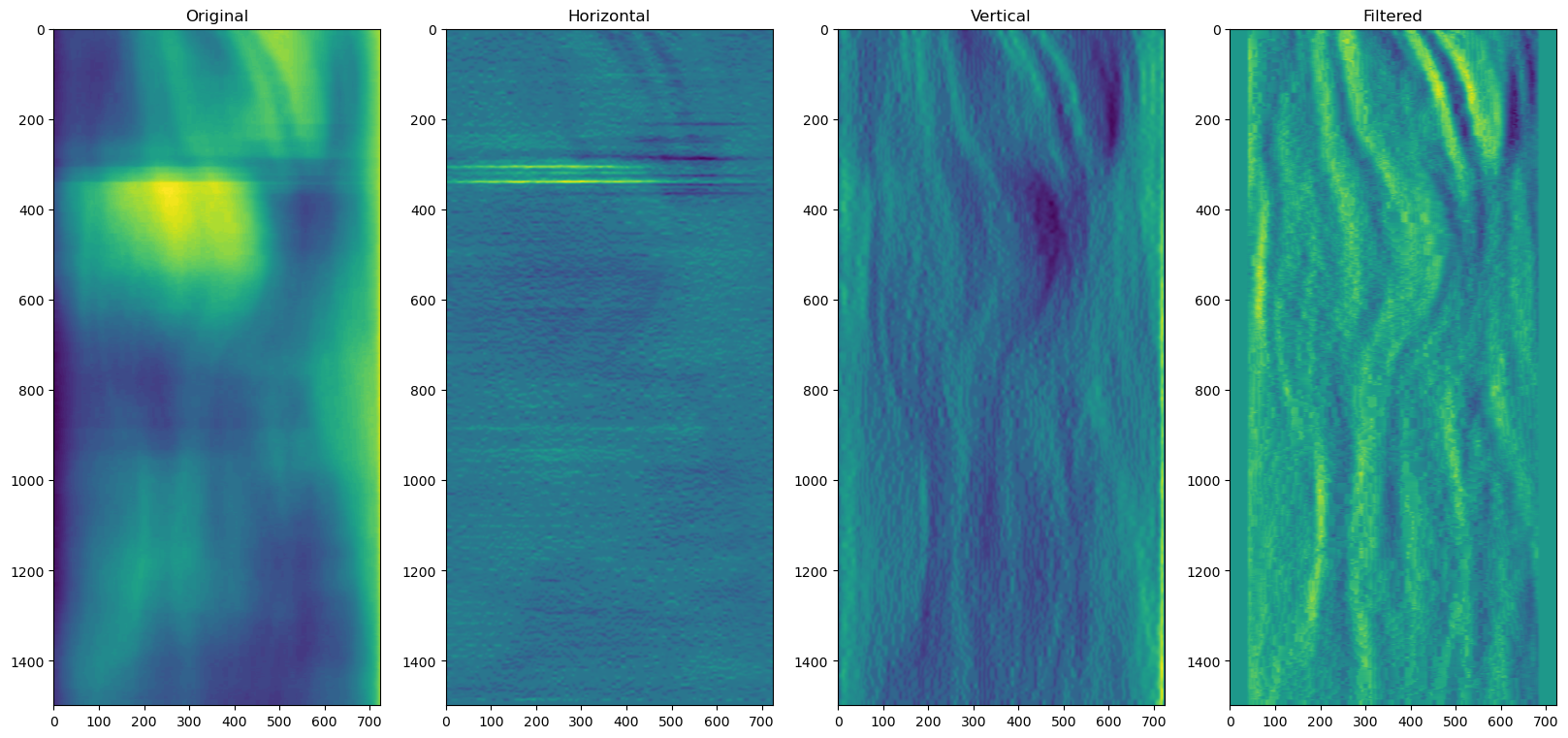

======================== Data Output==========================

A Mask Area Folder is created after running step 1 of the code. In this folder the mask area graphs have images with marked changes in intensity (blue (low intensity)/green (medium intensity)/yellow (high intensity)) and uterine horn borders (in red) from each frame of the .AVI video. These mask area files are used for constructing the area and mean intensity graphs in step 2 of the code.

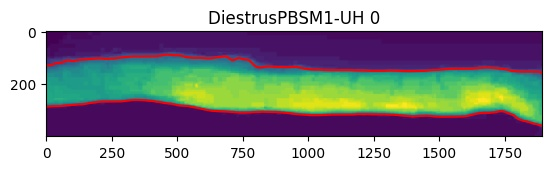

Area Graphs (jpg) and Mean intensity graphs (jpg)

Normalized graphs in a jpg format are generated. Wave coordinates cannot be extracted from these images so .html files need to be used (see next point).

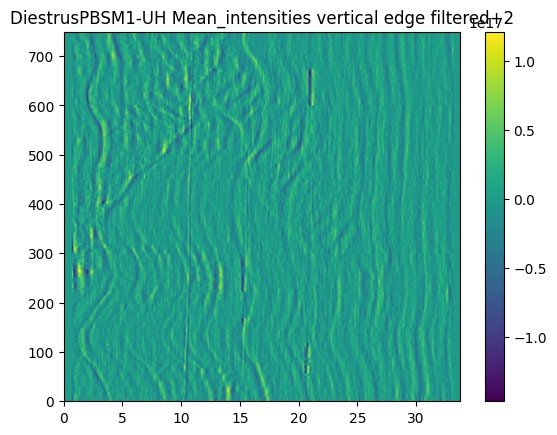

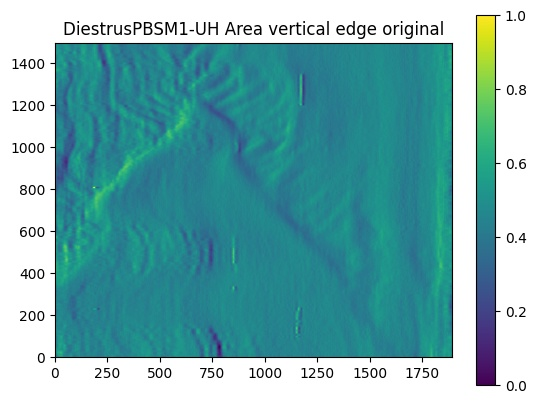

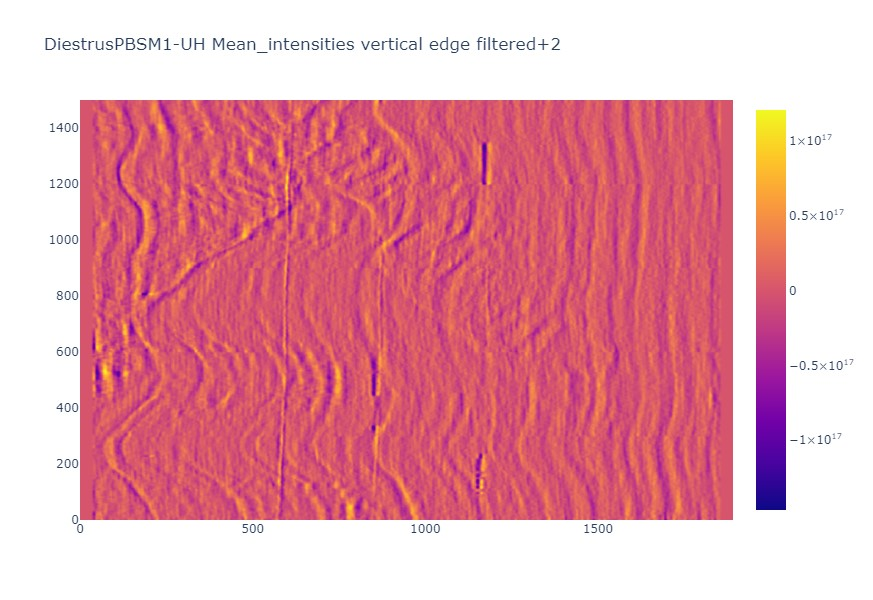
Area graphs or Mean Intensity Graphs (html):This output html file can be opened in a browser and wave coordinates are manually extracted by marking waves and calculating wave form metrics.

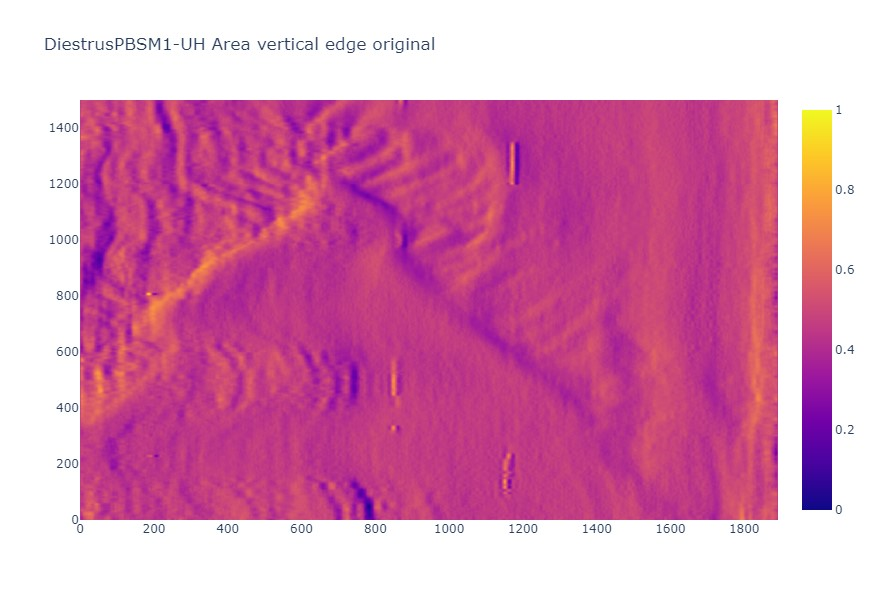
